# A unifying principle for multispecies coexistence under resource fluctuations

**DOI:** 10.1101/2024.10.28.620751

**Authors:** James A. Richardson, Jan Engelstädter, Andrew D. Letten

**Affiliations:** School of the Environment, The University of Queensland, Brisbane, Queensland 4072, Australia

## Abstract

Resource fluctuations are ubiquitous in nature and yet are generally assumed to play a limited role in the maintenance of biodiversity. We challenge this assumption by analyzing resource competition dynamics under conditions where traditional theory fails. We show that multi-species coexistence can be sustained when species are able to specialise on different temporal patterns of resource variability, including the asymmetries and periodic extremes commonly observed in natural systems. We further show how this partitioning of the statistical moments of the resource distribution provides a unified framework for explaining coexistence in variable resource environments. A theory of diversity maintenance based on moment partitioning highlights the potential for anthropogenic changes in resource regimes to drive cascading biodiversity losses.

## Main

Resource availability fluctuates through time in most natural systems. It is well known that such fluctuations can theoretically allow more than one species to coexist on a single resource *(1–3)*, in violation of ecology’s competitive exclusion principle *(4)*. Based on prevailing theory, however, it is generally assumed that resource fluctuations are inadequate to explain the large numbers of species commonly observed coexisting in nature *(2, 5–9)*. Less widely recognised are the restrictions imposed by the underlying theory, namely that: i) resource fluctuations need to be small in amplitude; and ii) asymmetries in the temporal distribution of resources are ignored. In real ecosystems, however, resource availability can fluctuate dramatically over multiple time-scales, with periodic extremes driven by diverse natural and anthropogenic processes *(10–16)*. As a result, we lack insight into competitive interactions under realistic resource regimes, and their implications for the maintenance of diversity.

A well-studied condition for coexistence in non-equilibrium environments is that species trade off competitive ability under variable versus continuous resource supply *(1–3)*. The diversity of terminology used to describe this dichotomy (and its spatial analogues) stands testament to its perceived ubiquity across diverse systems (gleaners and opportunists/exploiters *(17, 18)*, oligotrophs and copiotrophs *(19)*, competitors and colonizers *(20)*, digger and grazers *(21)*, cream-skimmers and crumb-pickers *(22)*). Alongside the trade-off itself, an additional condition for stable coexistence is that each species, when abundant, drives resource variability in a direction that favours its competitor *(1–3)*. Species favoured under constant resource supply need to increase resource variability, while species favoured under variable resource supply need to decrease resource variability. If these feedbacks are sufficient to yield negative frequency dependence, the outcome is coexistence. Levins *(23)* classically framed this trade-off as between competitors that consume the ‘mean’ of the resource and those that consume its ‘variance’.

Later work formalised the underlying mechanism of coexistence as ‘relative nonlinearity of competition’ (in reference to requisite differences in the nonlinearity of competitor growth curves) *(2, 24, 25)*, under the simplifying assumption of the so-called small variance approximation. This amounts to the requirement that the range of resource variation is sufficiently small that the growth curves of all competitors can be reasonably approximated by quadratic functions (akin to those studied by Levins *(23)*). This limits coexistence to just two species on a single fluctuating resource *(2, 23)*, an observation that has buttressed the widely held notion that resource variability is unlikely to be an important driver of diversity maintenance in nature *(2, 5–9)*.

In most natural systems, however, the small variance approximation is unlikely to hold. Natural cycles in weather, climate, circadian rhythms and patterns of feeding can drive dramatic fluctuations in nutrient availability to macro- and micro-organisms, on time-scales ranging from seconds to years *(26–28)*. These cyclic phenomena may in turn be perturbed to asymmetric extremes by a variety of processes including runoff, fires, glacial melt, coastal upwelling, drought, flooding, disease and dysbiosis *(10–16, 29)*. In spite of well-documented impacts on nutrient cycling and the frequency of extreme events under global change, the extent to which these more realistic patterns of resource variability affect the maintenance of biodiversity has received remarkably little attention.

To investigate the scope and generality of coexistence via resource fluctuations, we analysed a broad spectrum of resource competition dynamics under conditions where the small variance approximation fails. Building upon the work of Levins *(23)*, we introduce a technique for separating consumer growth rates into contributions that extend to the higher moments (beyond variance) of the resource distribution. When allowance is made for patterns of resource variability that capture the imbalances and periodic extremes of realistic systems, we discover a substantial increase in the bandwidth for multi-species coexistence. We further demonstrate how the principle of moment partitioning provides a common currency for explaining multi-species coexistence under resource fluctuations.

### An extended gleaner-opportunist trade-off

We model resource-consumer interactions using a standard formulation *(17, 30)*. To describe the dynamics of *n* species competing for a single resource, we employ a system of ordinary differential equations:

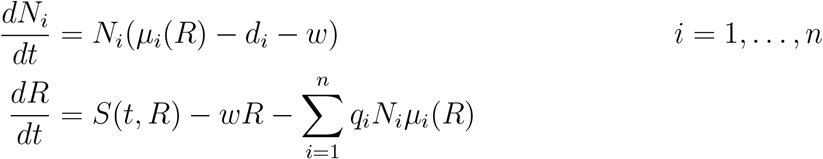

Here *N*_*i*_ is the population size of species *i*; *R* is the resource abundance; *μ*_*i*_ is the resource-dependent per capita growth rate of species *i*; *d*_*i*_ is its constant per capita mortality rate; *q*_*i*_ is the resource quota of species *i* (i.e. amount of resource consumed per unit biomass); *w* is the system washout rate (a constant rate at which all consumers and resources leave the system); and *S* is the resource supply function, which may vary with time and resource abundance. We consider various forms for both *μ*_*i*_ and *S*, allowing us to investigate both endogenous and exogenous sources of resource fluctuations. The general results are independent of the source of fluctuations.

First, we consider a three species system with 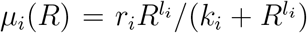 for each species. For *i* = 1, 2, we take *l*_*i*_ = 1, giving us Monod (i.e. Holling type II) functions with maximum per capita growth rate *r*_*i*_ and half-saturation constant *k*_*i*_. For *i* = 3, we take *l*_*i*_ = 2, giving a sigmoidal (type III) function (cf. previous studies, which have focused almost exclusively on consumers with type I or type II per capita growth rates *(1, 7, 31–33)*). Resources grow logistically with *S*(*R*) = *aR*(1 − *R/K*), where *a* is the resource’s intrinsic growth rate and *K* is its carrying capacity. The washout rate *w* is set to zero. We choose parameters (see supplementary materials section S1) so that species 1 and 2 exhibit the well-known gleaner-opportunist trade-off *(3, 17)*, giving the gleaner a higher growth rate at low resource abundance and the opportunist a higher growth rate at high resource abundance. The parameters for species 3 are chosen so that it has the lowest growth rate of the consumers at low resource abundance, and the highest at high resource abundance (Fig 1); since this is a magnification of the opportunist’s traits relative to the gleaner, we call species 3 a *supertunist*.

**Figure 1:**
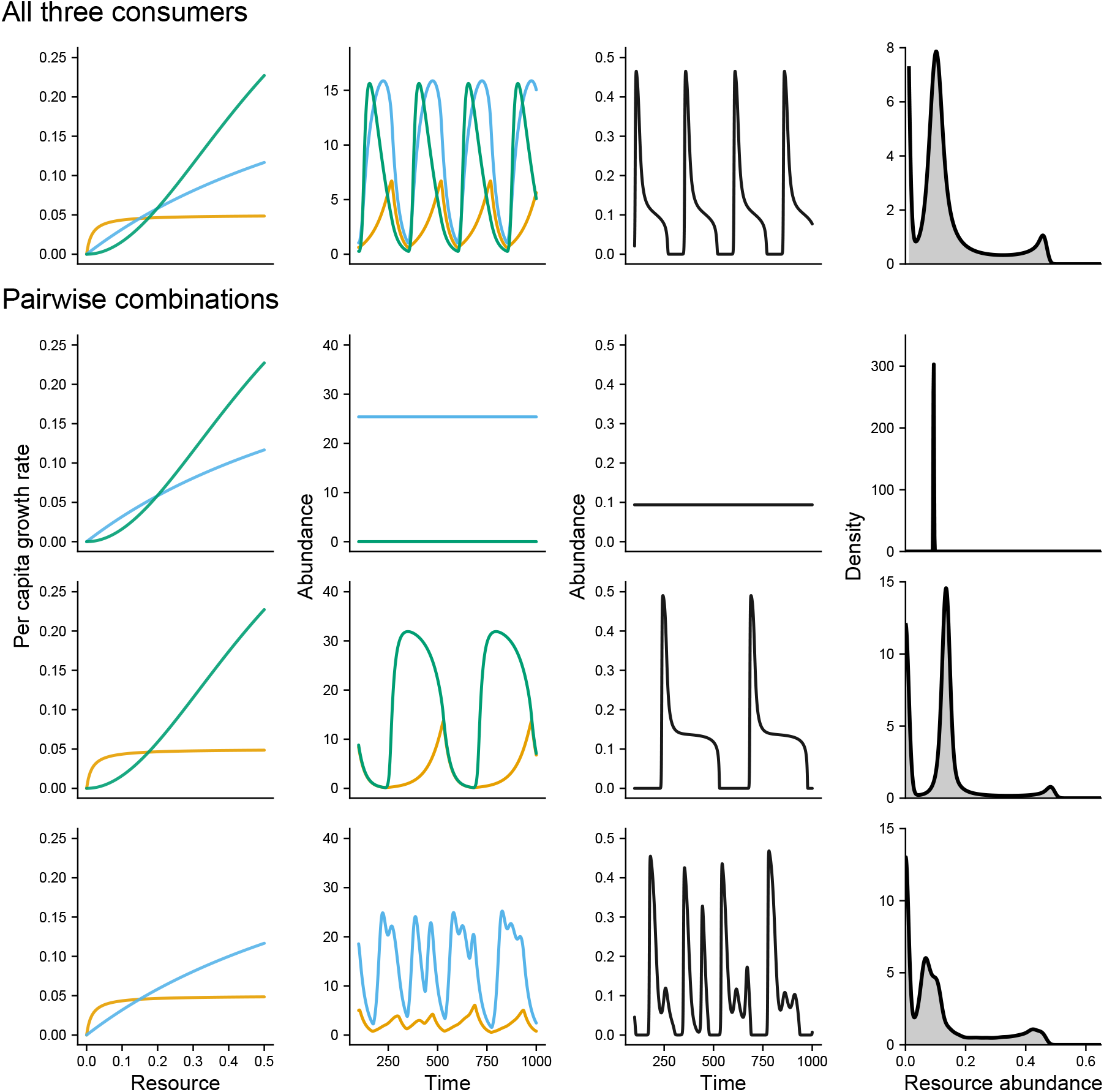
Coexistence in a gleaner-opportunist-supertunist system. First column shows the per capita growth rates of the gleaner (orange), opportunist (blue) and supertunist (green) in: i) the three species system (first row); ii) the opportunist-supertunist system (second row); iii) the gleaner-supertunist system (third row); and iv) the gleaner-oppertunist system (fourth row). Second column shows the corresponding steady-state population dynamics in simulations (after a burn-in of 5,000 time-steps). Third columns shows the corresponding resource dynamics over the same time period. Fourth column shows the corresponding kernel density estimates for the resource distribution through time. Without resource fluctuations introduced by the gleaner, the supertunist cannot survive alongside the opportunist. While the other systems approach a periodic orbit, the gleaner-opportunist system displays chaotic fluctuations. In each of the two-species systems, the resource distribution favours the invasion of the missing third species.

Contrary to theory based on quadratically approximated growth functions, which would limit coexistence under these conditions to two species, we observe stable coexistence of all three species in simulations (Fig 1). Comparison of each pairwise combination provides insight into the mechanism underlying coexistence. Because resource fluctuations are introduced endogenously by the gleaner, when the gleaner is absent, the resource is maintained at the opportunist’s *R*^*^ (equilibrium resource abundance when a single species is alone in the system) *(34, 35)*, and the supertunist is competitively excluded. In the gleaner-supertunist system, the supertunist responds rapidly to high resource abundance and quickly depletes the resource in the process. The supertunist’s population begins declining once the resource concentration drops below its *R*^*^, while the gleaner is able to maintain positive growth until the resource level collapses, at which point the gleaner’s own population collapses, the resource rebounds, and the cycle repeats (as in classic predator-prey cycling). The gleaner-opportunist system follows a similar pattern of booms and busts (mirroring the classic result of Armstrong and McGehee *(1)*), except the opportunist is able to maintain a positive growth rate at lower resource abundance, which leads to more peaks and troughs in the resource abundance through time, manifesting in greater extremes in the long-run resource distribution.

For stable three-species coexistence, each pair of species growing together must create conditions that benefit the third (Fig 1). When only the opportunist and supertunist are present, the system approaches equilibrium, which favours the gleaner with its low value of *R*^*^. When only the gleaner and supertunist are present, the resource distribution is concentrated around a medium value, which favours the opportunist, which has the highest per capita growth rate of the three species when resources are in this range. Similarly, when the gleaner and opportunist are the only species present, the more frequent peaks in resource abundance favour the supertunist, which has the highest per capita growth rate at high resource abundance. Moreover, since the supertunist is a poor competitor at medium resource concentrations, frequent drops to very low resource levels, where all three consumers grow almost equally poorly, are to its relative advantage. The resource distribution in the three-species system has features of the distributions from each of the two-species systems, reflecting the combined effects of all three species. When an absent species enters any of the two-species systems, it makes the resource distribution less favourable to itself. This is a hallmark of the negative frequency dependence that underpins stable coexistence. We show that this system remains stable for a wide range of resource parameters in the supplementary materials section S2.1 and fig S1. Note that although stable persistence of three type II competitors is also technically possible (supplementary materials section S3.1, figs S5, S6), previous research has found limited bandwidth for coexistence *(8, 9)*).

### Coexistence through moment partitioning

We now discuss a more analytic approach to understanding stable coexistence under resource fluctuations. If species *i* is to persist in the system without going extinct, then the long-term average of its per capita growth rate must balance its per capita rate of loss from the system.

This gives us the following equality, formally derived by Levins *(23)*:

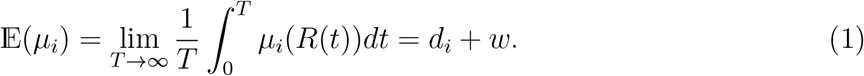

In addition to this persistence condition, the standard invasion condition tells us that species *i* can invade a system from low numbers if

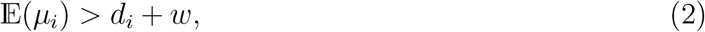

where this time the average is calculated with species *i* absent from the system. For all species to stably coexist, Conditions 1 and 2 must be met for every species.

We illustrate these conditions with a three-species system, with per capita growth rates all given by polynomials. This example extends the model described by Levins *(23)* to include a consumer with a cubic per capita growth rate.

We consider per capita growth rates of the following form:

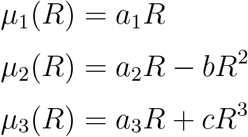

Resource fluctuations are supplied exogenously (see supplementary materials section S4 for details), which provides greater control over resource variability than endogenous fluctuations, and thereby allows us to create an illustrative example where the desired behaviour can be observed.

Taking averages, we can express the average per capita growth rates in terms of the moments of the resource distribution:

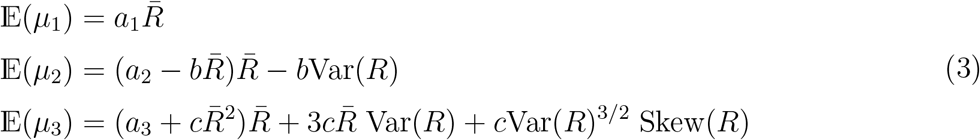

Here 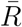 is the average resource abundance in the system, Var(*R*) is the variance of the resource distribution and Skew 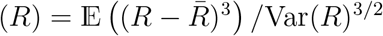 is its skewness.

We choose parameters (supplementary materials section S4) such that species 1 (linear), 2 (quadratic) and 3 (cubic) respectively approximate the opportunist, gleaner and supertunist strategies of the previous example (Fig 2A). Using Condition 1, when all three species are coexisting, the system of equations (3) determines the mean, variance and skewness of the resource distribution. Using these equations we can directly compare the contribution each moment makes to each species’ average growth rate at equilibrium (Fig 2B).

**Figure 2:**
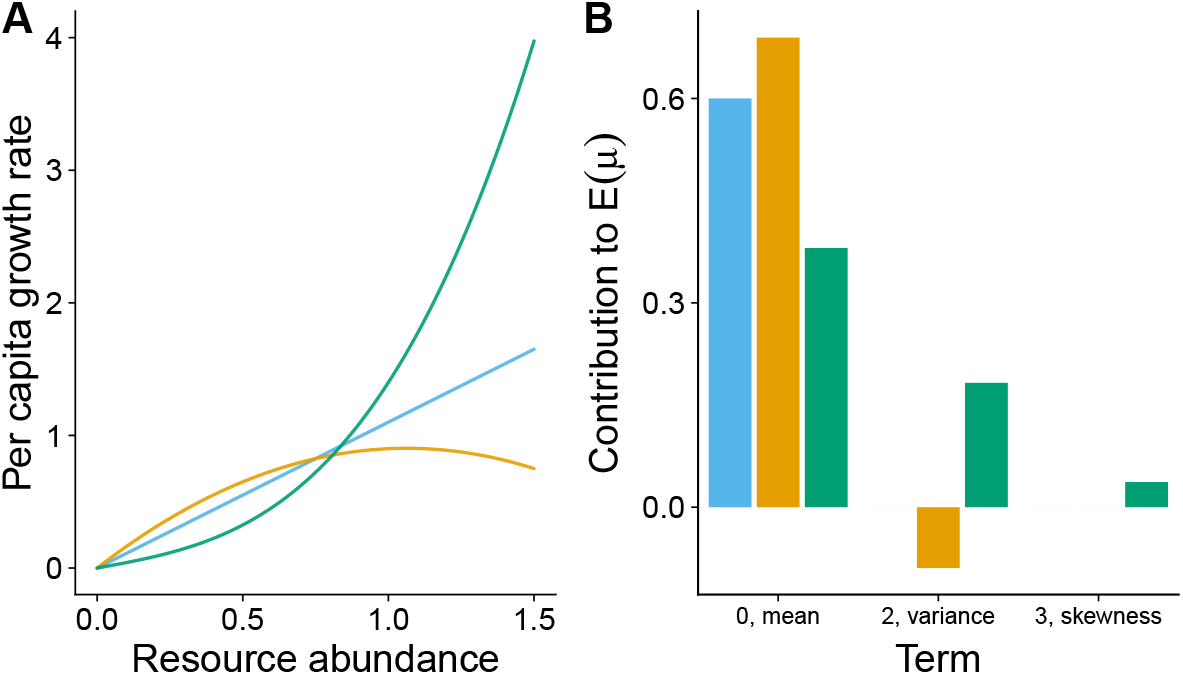
A three species system with polynomial per capita growth rates. **A**. Per capita growth rates of the opportunist (linear, blue), gleaner (quadratic, orange) and the supertunist (cubic, green). High resource abundance inhibits growth of the gleaner. **B**. Each term in the system of equations (3), calculated in the three-species system. Bars represent terms in the expression for the average per capita growth rate of the gleaner (orange), opportunist (blue), and the supertunist (green). The zeroth term corresponds to the average, the second term to the variance, and the third to the skewness. Note that the first term vanishes (cf. Fig 3D and fig S8). By Condition 1, each expression sums to *d*_*i*_ + *w* = 0.6 for each species. From **B**, we can read off the contribution each moment makes to each consumer’s average per capita growth rate in the three-species system.

We use the invasion condition to describe one possible community assembly pathway (see supplementary materials section S4 for details). The linear opportunist can enter the system if the mean resource abundance is sufficiently high. Once in the system, by Condition 1, the opportunist determines the mean resource abundance. In this way, the opportunist can be said to consume the mean of the resource distribution *sensu* Levins *(23)*. With the opportunist already in the system, by Condition 2, the gleaner can invade if the variance in the resource distribution is sufficiently small; its quadratic per capita growth function results in negative growth when resources deviate too far either side of the mean (this differs from Levins’ *(23)* scenario, where the quadratic competitor benefits from variance and ‘consumes’ it in the process). In contrast with the quadratic gleaner, the cubic supertunist can invade if the variance and skewness combine to be sufficiently large, reflecting the fact that the supertunist benefits especially from significant deviations above the mean (Fig 2A). We can use simulations to see that the variance and skewness of the resource distribution in the opportunist monoculture satisfies the invasion condition for the supertunist but not the gleaner. However, once the supertunist invades the opportunist monoculture to form a two species system, the variance is reduced sufficiently for the gleaner to invade and establish the complete three-species system (see fig S9). In summary, the opportunist, gleaner and supertunist occupy unique niches defined by the mean, the variance and the skewness, respectively.

### Beyond polynomials

We now consider how to use Conditions 1 (persistence) and 2 (invasion) to express how different moments of the resource distribution benefit or harm a consumer’s growth in the more practical case where the consumer’s per capita growth rate *μ*(*R*) is described by a conventional functional form (i.e. non-polynomial; we focus on type II and III functional forms, but the results apply generally). Suppose *μ*(*R*) admits a Taylor expansion around the point *R* = *Z*. Then we can calculate 𝔼 (*μ*) term-by-term using the Taylor expansion (see supplementary materials section S6 for details). We explicitly describe the procedure below in the case that *μ*(*R*) is type II. If we take 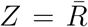 (average resource abundance), this procedure returns the average per capita growth rate of species *i* in terms of the familiar central moments of the long-term resource distribution. This mirrors the example given above for consumers with polynomial per capita growth rates. However, we illustrate below that other choices of *Z* may also be necessary and informative.

Given a type II function, its Taylor expansion around *R* = *Z* is as follows (see derivation in supplementary materials section S5):

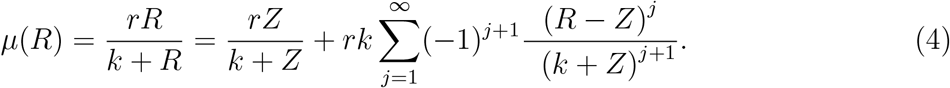

This converges for *R < k* + 2*Z*. Thus, if resources are bounded strictly below *k* + 2*Z*, we obtain an expression for 𝔼 (*μ*) in terms of moments of the resource distribution, by taking the average within the sum (supplementary materials section S6):

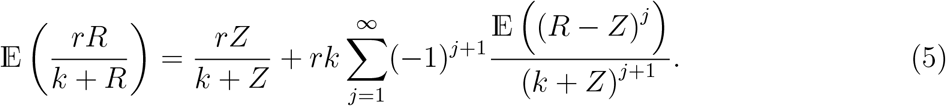

If we take 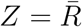, the first few terms of Equation 5 are as follows:

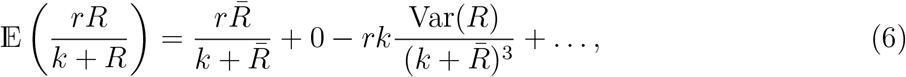

This expression for 𝔼 (*μ*) in terms of moments can be substituted into Conditions 1 and 2 to understand how species in a given system can coexist.

### Relative nonlinearity and the small-variance approximation

This approach of using Taylor expansions to approximate non-polynomial per capita growth rates is similar to the approach taken in coexistence theory *(2, 25)*. A critical difference, however, is that the machinery of coexistence theory requires that we truncate the Taylor expansion at order 2 to obtain a quadratic approximation. To make this possible, the variance in the resource distribution must be sufficiently small to ensure the approximated per capita growth function does not deviate too far from the true function. If this small variance approximation *(36)* is not valid, then the classical methods of coexistence theory cannot be applied, and much of the intuition inherited from coexistence theory fails.

In a classical gleaner-opportunist system *(1, 17)*, we find that the small variance approximation can only be satisfied if the two consumers possess improbably similar per capita growth rate functions, which severely restricts the system’s coexistence bandwidth (see supplementary materials section S7 and fig S10). To achieve a larger coexistence bandwidth, Abrams and Holt *(8)* show that the gleaner’s per capita growth rate must be more saturating, which corresponds to a larger number of non-negligible terms in Equation 6, violating the small variance approximation. Thus, even in these classical systems, the small variance approximation is unlikely to hold if the system displays any kind of robust coexistence. Despite this limitation, the special properties of systems that satisfy the small variance approximation have helped perpetuate the impression that resource fluctuations are an ineffective driver of coexistence. In the next sections, we show that opportunities for species coexistence via relative nonlinearity expand significantly when the constraints of the small variance approximation are relaxed.

### Beyond the small-variance approximation

We now illustrate how the Taylor expansion can be used to isolate the contributions that the higher moments (beyond variance) of the resource distribution make to species invasion and persistence. Note that even moments, like variance, only capture the magnitude of deviations around the mean, while odd moments, like skewness, also capture their asymmetry. Consider first a stable system comprising a gleaner and an opportunist, both with type II per capita growth rates (Fig 3A&B). Resource fluctuations are supplied exogenously via continuous sinusoidal variations in resource supply (see supplementary materials section S1 for full model details; qualitatively identical results can be obtained by modelling resource supply in discrete pulses [section S9]). When the gleaner is alone (shown in orange in Fig 3C), resource abundance varies more widely. The opportunist benefits, relative to the gleaner, from this increased variation, and is thus able to invade the gleaner monoculture. Conversely, the opportunist creates conditions with less variation that benefit the gleaner.

**Figure 3:**
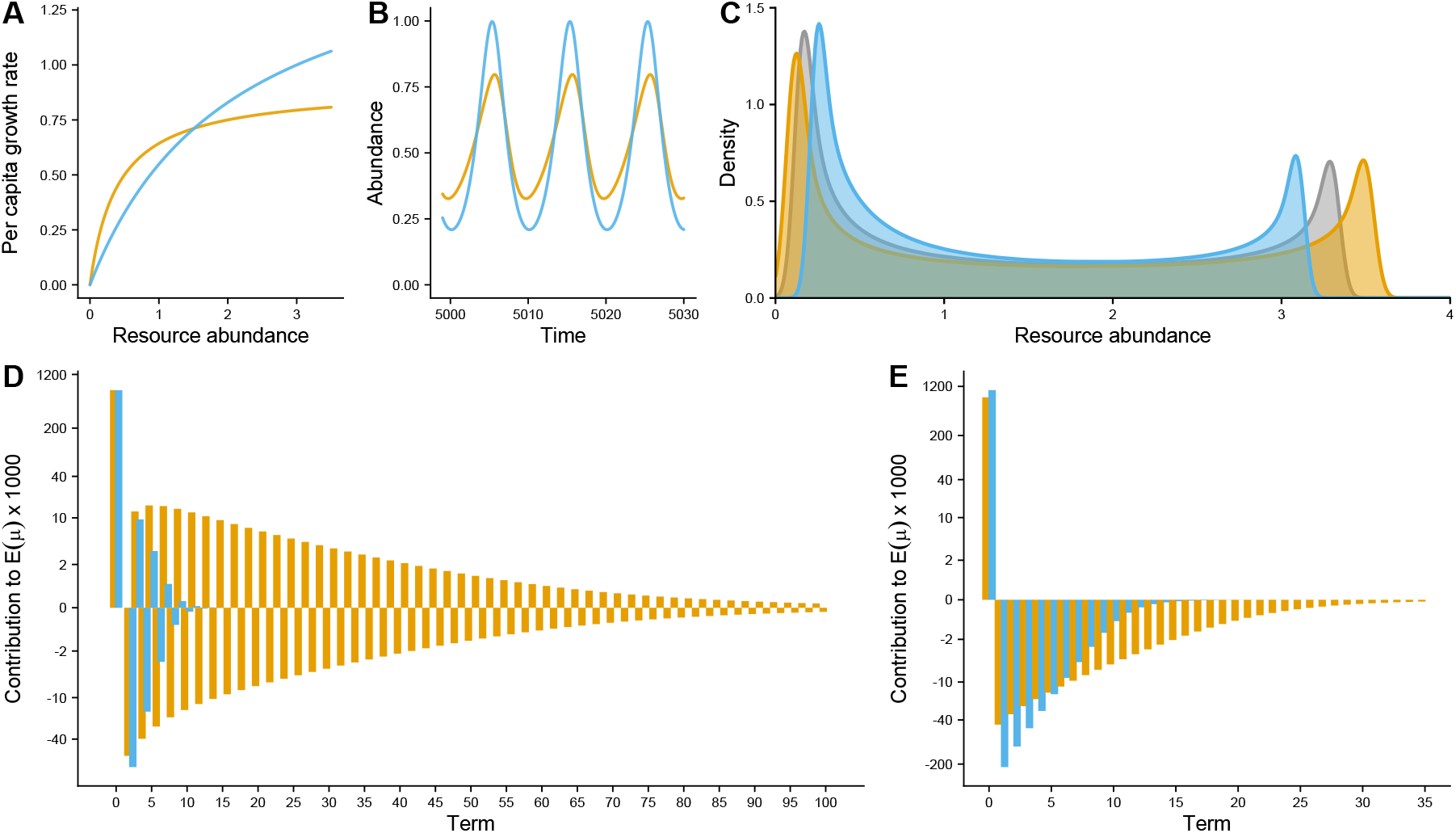
Moment partitioning in a gleaner-opportunist system. **A**. Per capita growth rates of the opportunist (blue) and the gleaner (orange). **B**. Simulation results showing steady-state population dynamics. **C**. Kernel density estimates for resource abundance through time. Variation in the resource distribution is greatest in the gleaner monoculture (orange), and least in the opportunist monoculture (blue), which in either case benefits the absent species. Resource variation is intermediate in the two-species system (grey). **D & E**. Terms in the expansions of each consumer’s growth rate around 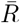 (**D**) and *R*_max_ (**E**) in the two-species system. Average per capita growth rates calculated from simulations of the two-species system (term contributions sum to *d*_*i*_ + *w* = 0.6 but multiplied by 1000 for visualization). Although 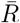 and *R*_max_ change in the species monocultures, the qualitative shape of the plots remains the same (see fig S11). Parameters listed in the supplementary materials section S1.

To see how the the gleaner and opportunist partition the moments of the resource distribution, we expand the per capita growth rates of each species around 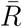, as in Equation 6 (Fig 3D shows the size of each term). It is evident that higher moments make a non-negligible contribution to the average per capita growth rate of each species, indicating the failure of the small variance approximation. Moreover, the opportunist is more negatively affected by variance than the gleaner, an observation that cautions against the dominant narrative that opportunist strategists are the primary beneficiary of resource variance *(3, 25)*. Nevertheless, the combined effect of all higher moments on the opportunist is still less negative (fig S12). Not only does the opportunist benefit from resource deviations above the mean, it also benefits relative to the gleaner from large deviations below the mean, which erode the gleaner population and allow the resource to rebound. As such, the opportunist can still be said to benefit from higher moments in the resource distribution relative to the gleaner.

Rather than expand around 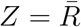, we can present the same information in terms of moments centred around *Z* = *R*_max_, the maximum resource value (Fig 3E). Substituting *Z* = *R*_max_ into Equation 5, the first term in the expansion records the focal species’ growth rate at the maximum resource concentration. All subsequent terms record the effect of deviations below the maximum (as in the familiar case of moments centred around the mean, higher moments give greater weight to larger deviations). Since the only possible deviations around the maximum are negative, odd higher moments (that is, 𝔼 ((*R* − *R*_max_)^*j*^) for *j* odd) are now always negative, making all but the first term in Equation 5 also negative. While the opportunist grows better than the gleaner at *R*_max_, small deviations below the maximum are more detrimental to the opportunist than the gleaner, since the opportunist’s per capita growth rate declines more steeply than the gleaner’s as resource abundance drops below the maximum. Thus, the opportunist and the gleaner are more negatively affected by smaller and larger moments, respectively.

We now return to the three-species system illustrated in Fig 1. The expansions around the average do not converge in this system, so we expand around *Z* = *K*, the carrying capacity of the logistic resource (Fig 4; see supplementary materials section S5 for a derivation of the supertunist’s type III Taylor expansion). This choice has the advantage of being the same across the two-species systems and the complete three-species system, making for easy comparison of the expressions we obtain for the invasion and resident average per capita growth rates. Not only does the supertunist benefit from resources being close to carrying capacity, but higher moments, which reflect large deviations below the carrying capacity, also benefit the supertunist relative to the other consumers (Fig 4). Smaller moments, corresponding to relatively small drops below the carrying capacity, are more detrimental to the supertunist than its competitors, reflecting the steep decline in its per capita growth rate at high resource levels. With both the gleaner and the opportunist exhibiting a similar distribution of terms as in the pairwise scenario described above (Fig 3E), it is clear that the three species access different moments in the resource distribution. This partitioning of the moments explains their coexistence.

**Figure 4:**
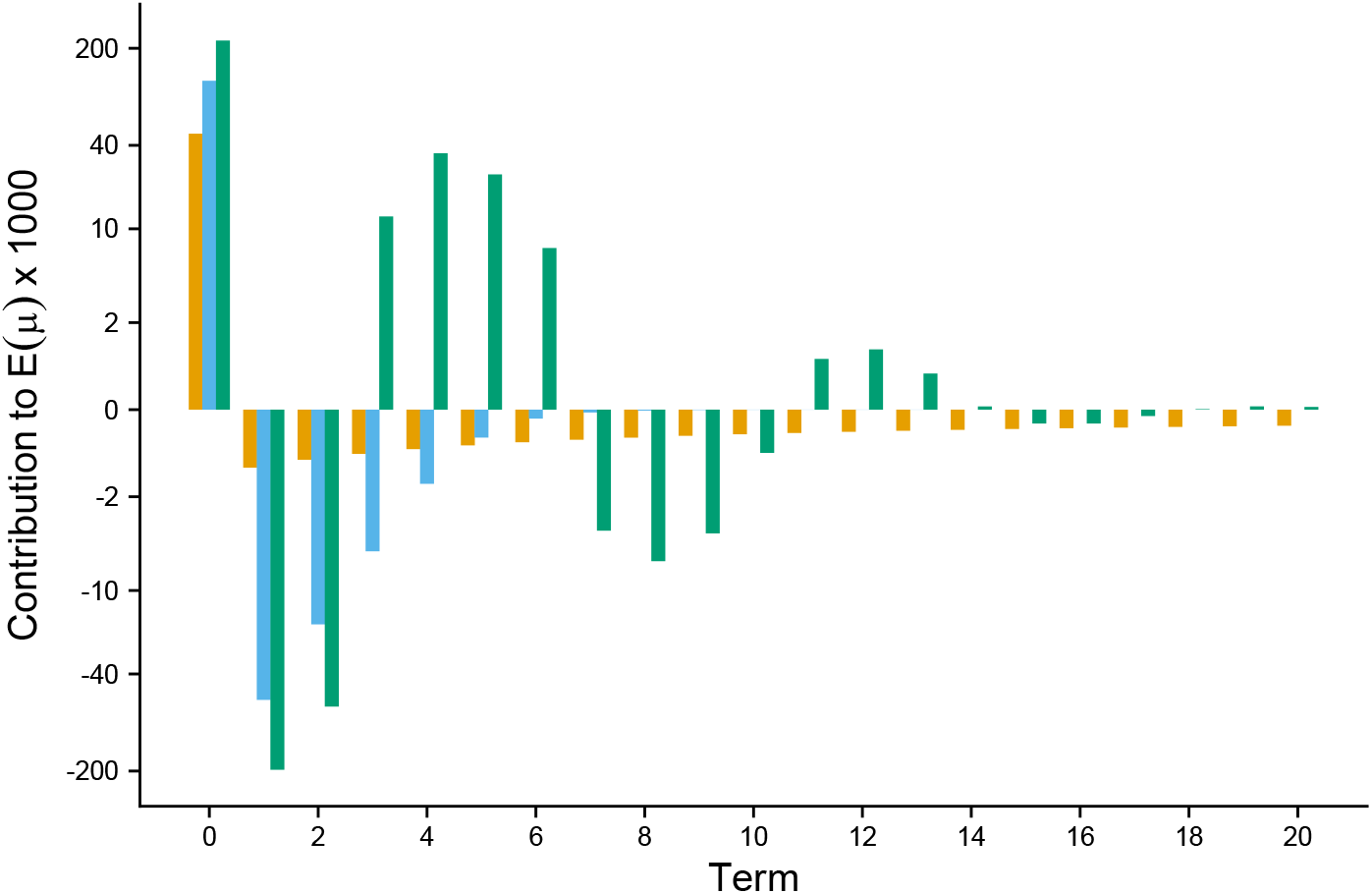
Moment partitioning in a gleaner-opportunist-supertunist system. The terms in the expansions of the gleaner’s (orange), the opportunist’s (blue) and the supertunist’s (green) average per capita growth rate around the resource carrying capacity *K* (*R*_max_). Term contributions sum to the consumer’s average per capita growth rate, which is equal to the common death rate *d*_*i*_ = 0.03 (multiplied by 1000 for visualization). Of the three consumers, the gleaner grows slowest when *R* = *K* (term 0), but low-order moments, which capture relatively small deviations of the resource below *K*, have a smaller negative effect on the gleaner than on its competitors, since in this region the gleaner’s per capita growth rate curve is close to constant (Fig 1). In contrast, higher-order moments, which capture large deviations below *K*, have a larger negative effect, reflecting the steep decline in the gleaner’s per capita growth rate when resources are scarce. The opportunist is inhibited more than the gleaner by low-order moments since its per capita growth rate declines more steeply at resource levels near *K*, but the effect of high-order moments is negligible. The supertunist suffers most from low-order moments because it has the steepest per capita growth rate near *K*. However, since its per capita growth rate levels out at low resource abundance, the contribution of higher-order moments to the supertunist’s average per capita growth rate partially cancels out the large negative contribution of the lower-order moments.

Although we define the supertunist in an explicit functional form (i.e. type III), it should be noted that type-III-like and other non-canonical per capita growth responses can also emerge through a range of more complex and/or adaptive organismal behaviour, such as rapid evolution *(3)*, access to dormant life stages *(37)* or resource storage *(29, 38)*. In many cases, we should expect the relative nonlinearities of these emergent growth responses to be sufficiently differentiated for moment partitioning to arise. Nevertheless, the reverse is also possible, where ostensibly different life histories converge on the same niche. For example, we can replace the supertunist by a species, which we term a survivalist, which employs a strategy similar to the dormancy strategist considered in *(37)*. While its strategy is ostensibly ecologically distinct from the supertunist’s, the survivalist benefits from similar moments in the resource distribution (supplementary materials section S10; fig S20), and therefore they occupy similar niches. If all four species are initiated in the system, the supertunist is driven to extinction.

### Many-species coexistence

Provided their per capita growth rate functions are sufficiently differentiated, coexistence via moment partitioning extends to systems with higher numbers of consumers. Fig 5 shows a system with five consumers, coexisting on a single fluctuating resource, where the unique consumer strategies are apparent in their different population dynamics through time (Fig 5B). As in Fig 1, consumers with higher maximum growth rates grow faster initially but peak earlier. The differing extent to which each consumer benefits from extremes in the resource distribution allows all five to coexist.

**Figure 5:**
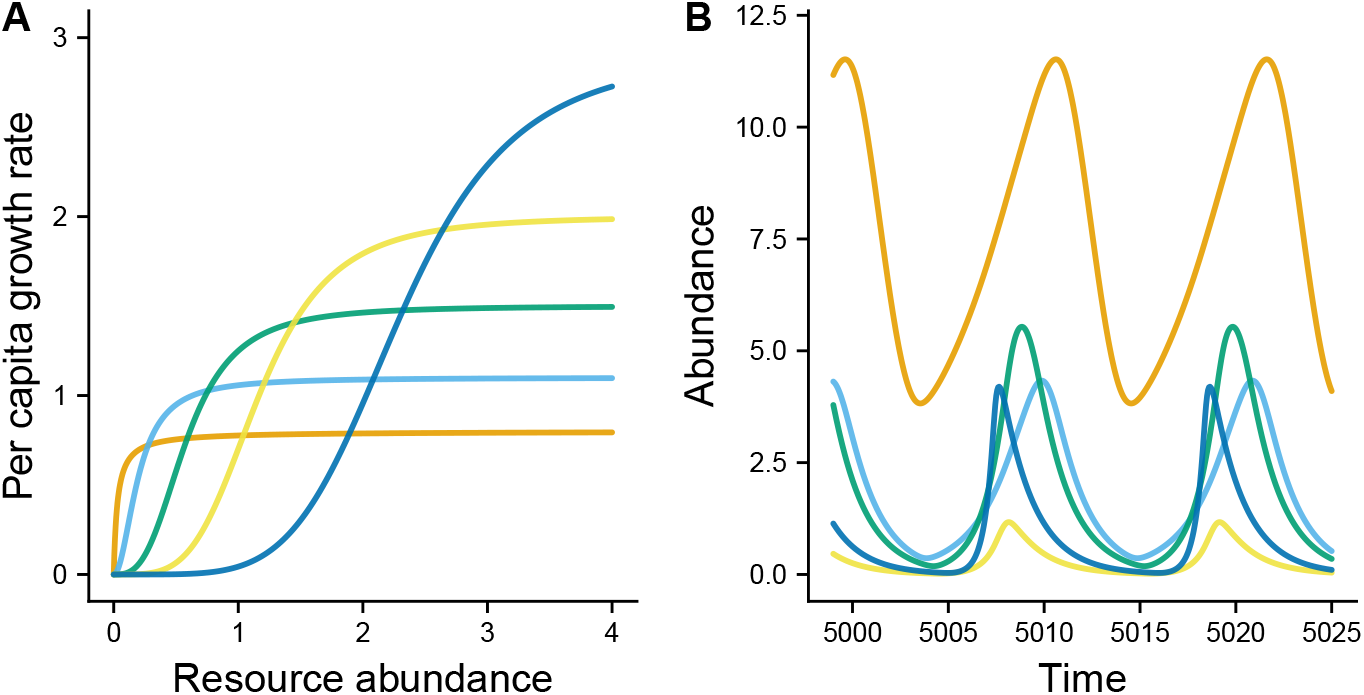
Coexistence of five-species on a single resource. **A**. Per capita growth rates of each species. **B**. Simulation results showing steady-state population dynamics. Consumer populations peak one after the other within the space of a single resource cycle, reflecting the different consumer strategies. Parameters listed in the supplementary materials section S1.

A notable feature of this system is that it is characterised by emergent coexistence *(39, 40)*, whereby multi-species communities are able to coexist even when many of the constituent species cannot persist in smaller assemblages (e.g. pairwise). For example, if the gleaner is absent, the system collapses to a single species (the species with the second lowest *R*^*^ value, shown in light blue in Fig 5). Several of the systems we investigated display similar behaviour (supplementary materials section S11), reflecting the emergence of new niches through changes in the resource distribution brought about by consumer-resource dynamics. In line with Levins’ analogy that species consume moments in the resource distribution *(23)*, we can interpret this phenomenon as a form of moment ‘cross-feeding’.

## Discussion

Resource fluctuations are well known to support coexistence in models of simple two-species communities *(1–3)*, but the prevailing view has long been that they cannot account for the diversity found in real-world systems. *(2, 5–9)*. Our findings suggest that this conclusion is grounded in an overly simplistic binary classification of species into equilibrium versus non-equilibrium resource specialists (e.g., gleaners and opportunists). In real systems, a wide variety of endogenous and exogenous forces cause resources to fluctuate over multiple timescales, presenting species with diverse temporal niches for exploitation *(26–28, 38)*. When allowance is made for the periodic extremes and asymmetries of natural systems, we find, contrary to previous work, that multiple species can in fact persist on a single fluctuating resource.

We propose that the vast majority of ecosystems exhibit ample temporal complexity to allow moment partitioning to operate *(26)*. Some familiar examples of organisms adapted to the higher moments of resource distributions include blue-green algae (e.g. *Nodularia, Dolichospermum* and *Microsystis* spp.) that thrive following the extreme nutrient pulses accompanying oceanic upwelling or agricultural runoff into lakes and rivers *(41)*, and plants that dominate in nutrient-rich post-fire environments *(42)* or that only emerge in response to rare, heavy rainfall such as desert annuals *(43)*. On shorter time-scales, the periods of feast and famine experienced by bacteria in the animal gut foster coexistence of distinct taxa that peak at different times post feeding *(44, 45)*. As such, a better understanding of diversity maintenance via moment partitioning is relevant across fields - from conservation and environmental management to agriculture and human health - particularly in light of ongoing anthropogenic changes to resource regimes *(46–48)*.

When resource fluctuations are not artificially constrained, a handful of previous studies have hinted at the broader potential for multi-species coexistence on a single resource *(29, 37, 38, 49, 50)*. In particular, several authors have reported opportunities for robust multi-species coexistence when consumers exhibit complex behaviour such as resource storage and dormancy *(29, 37, 38)*. The unfortunate caveat of this earlier work has been the need to appeal to the specifics of a particular system and set of ecological strategies. Our results provide a general framework for understanding multi-species coexistence under resource fluctuations, independent of the underlying biological mechanism.

Notwithstanding the non-canonical growth responses that likely arise from adaptive behaviour and other complex consumer-resource dependence, we emphasize that type III responses are sufficient to substantially broaden the bandwidth for fluctuation-dependent coexistence compared to the well-studied case when consumer responses are limited to types I and II *(1, 7, 31, 32)*. While a considerable theoretical basis underpins the use of type III responses *(37, 51–53)*, they remain remarkably understudied in empirical research. This gap stems largely from the inherent challenge of obtaining sufficiently high resolution observations of population growth dynamics in most plants and animals *(53)*. The fast generation times and high-throughput of microbial systems likely offer the best prospect for robust empirical fits of type III responses, as well as for experimentally investigating the principle of moment partitioning. Note that reliable estimates of the higher moments require increasing sample sizes, owing to the larger weighting of outliers (an empirical limitation that might also explain why the higher moments have been overlooked in coexistence research).

Alongside experimental investigations, various opportunities exist to broaden the theory of moment partitioning. In the many systems with multiple resources, their covariances and higher-order cross-moments provide consumers with additional potential temporal niches *(23, 54)*. Although Levins alluded to this possibility *(23)*, to our knowledge this phenomenon has never been formally analysed (but see Huisman and Weissing *(55)*, in which specialisation on cross-moments likely contributes to multispecies coexistence). The methods introduced here can be extended to characterise the contributions of these cross-moments to coexistence. Similarly, the approach could be adapted to quantify diversity maintenance through resource variation in space as well as time. Finally, the neglected role of resource fluctuations in emergent coexistence represents a significant opportunity for further investigation, with important implications for ecosystem management at both micro and macro scales.

## Author contributions

Conceptualization: J.A.R. and A.D.L. Formal analysis: J.A.R. Supervision: J.E. and A.D.L. Visualization: J.A.R. and A.D.L. Writing – original draft: J.A.R. and A.D.L. Writing – review & editing: J.A.R., J.E. and A.D.L.

## Data and materials availability

Code for reproducing all analyses will be provided on GitHub and Zenodo at time of publication.

## Supplementary materials

Materials and Methods

Supplementary Text

Figs. S1 to S24

References *(56-61)*

## Materials and Methods

All simulations were run in R version 4.3.0 *(56)* using the lsoda solver in deSolve *(57)*. Simulations were initiated with each species’ population at 1. Depending on the resource supply regime, resource abundance was initiated as follows: at the carrying capacity *K* in the case of logistic supply, at 1 in the case of periodic continuous supply, and at the pulse size *S*_0_ in the case of discrete supply.

Simulations were considered to have reached a stationary cycle when the difference in each state variable from one period to the next was less than 0.0001. (Note that the period of the cycle is determined by the resource supply in the case of externally driven fluctuations, and was calculated numerically in the case of logistic resource supply.) All but one of the systems described across the main text and the supplementary information were run for 10,000 time steps, at the end of which all systems with stable limit cycles had reached stationarity. The only system that required longer to reach stationarity was the gleaner-opportunist-survivalist system shown in fig S21, which was run for 30,000 time steps.

All but one of the simulations we considered ultimately reached a stationary cycle. In such systems, we considered that a species was extinct if, over the final cycle of the simulation, its population never rose above 0.0001. Any species whose population reached levels higher than 0.0001 over the final period was considered to be coexisting in the simulation. This conclusion was confirmed using an invasion analysis, in which, for each species, the system was simulated with the same initial conditions except that that species’ initial population was set to zero. The average per capita growth rate of the absent species over the final cycle of the simulation was then calculated numerically. If the average per capita growth rate thus obtained was greater than the death rate, the species was considered to be able to invade. The only simulation that did not reach a stationary cycle was the gleaner-opportunist subsystem within the gleaner-opportunist-supertunist system of Fig 1, which underwent chaotic oscillations. (Note that the identical species pair also appears in the gleaner-opportunist-survivalist system.) To decide invasion from this system, the average per capita growth rate of the missing species was calculated numerically over the last 1500 time steps of the simulation.

Results of the invasion analysis agreed exactly with the results of the long-run simulations. Note that, for systems with three species or fewer, invasion analysis is always sufficient to determine coexistence, as long as the system does not display alternate stable states. This is not the case, however, for systems with more than three species. For a discussion of invasion in the five species system shown in Fig 5, see *S11 Community assembly*.

## Supplementary Text

### S1 Full models and parameter values from the main text

Here we provide the full details and parameters of all models from the main text, other than the model with polynomial per capita growth rates, which is discussed separately in *S4 An example with polynomial per capita growth rates*. All models have the following general form:

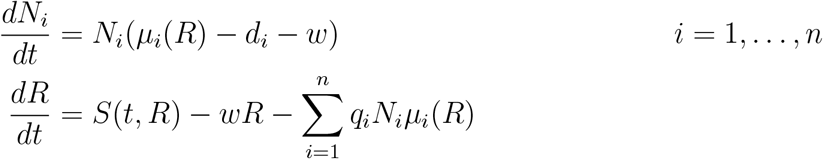

The gleaner-opportunist-supertunist system shown in Fig 1 has 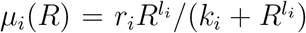 and *S*(*R*) = *aR*(1 − *R/K*). The parameters are *r*_1_ = 0.05, *k*_1_ = 0.015, *l*_1_ = 1, *r*_2_ = 0.35, *k*_2_ = 1, *l*_2_ = 1, *r*_3_ = 0.5, *k*_3_ = 0.3, *l*_3_ = 2, *d*_*i*_ = 0.03 for all *i, q*_*i*_ = 0.1 for all *i, a* = 1, *K* = 0.5, *w* = 0.

The gleaner-opportunist system shown in Fig 3 has 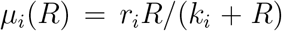 given by Monod functions, and *S*(*t*) = *A*(*α* sin(2*πt/P*) + 1). The parameters are *r*_1_ = 0.9, *k*_1_ = 0.4, *r*_2_ = 1.7, *k*_2_ = 2.1, *d*_*i*_ = 0.1 for all *i, q*_*i*_ = 2 for all *i, A* = 2, *α* = 0.8, *P* = 10, *w* = 0.5.

The five-species system shown in Fig 5 has 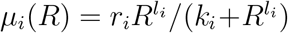 and *S*(*t*) = *A*(*α* sin(2*πt/P*)+ 1). The parameters are *r*_1_ = 0.8, *k*_1_ = 0.03, *l*_1_ = 1, *r*_2_ = 1.1, *k*_2_ = 0.04, *l*_2_ = 2, *r*_3_ = 1.5, *k*_3_ = 0.2, *l*_3_ = 3, *r*_4_ = 2, *k*_4_ = 1.85, *l*_4_ = 4, *r*_5_ = 2.9, *k*_5_ = 65, *l*_5_ = 5, *d*_*i*_ = 0.1 for all *i, q*_*i*_ = 1 for all *i, A* = 8, *α* = 0.9, *P* = 11, *w* = 0.5.

### S2 Robustness under parameter changes

For several of the systems presented in the main text, we used simulations to investigate their behaviour over a range of parameter values. These simulations illustrate the robustness of these systems under parameter change. In particular, the results of these simulations show that the many-species coexistence illustrated in the text is achievable over a range of different resource supply parameters.

In each case, we identified community persistence using invasion tests and long-run simulations. If the systems exhibit alternate stable states then this approach is susceptible to errors; however, using simulations to explore a range of initial conditions has not yielded any evidence of alternate stable states, and we are confident they are not exhibited in the any of the examples investigated.

#### S2.1 gleaner-opportunist-supertunist system

Figure S1 shows the community composition in the gleaner-opportunist-supertunist system shown in Fig 1 for a range of resource parameters. We have varied the resource carrying capacity *K* and its intrinsic growth rate *a*. Each simulation was run for 10, 000 time steps. If, over the last five time steps of the simulation, a species’ population was ever greater than 0.001, that species was considered to be persisting; otherwise it was considered to be extinct. Since the choices of cutoff are arbitrary and persisting species may be overlooked, plots of the simulation outcomes were checked manually to verify the behaviour around the boundary between the three-species and two-species coexistence regions. The approximate location of the three-species coexistence region was also confirmed using invasion analysis.

#### S2.2 Three species with type II/Monod per capita growth responses

Figure S2 shows the community composition over a range of resource supply parameters for the system discussed in *S3 Three consumers with type II (Monod) per capita growth rates*. Across all parameter combinations investigated, the gleaner species (lowest *R*^*^) always persists. Figure S3 shows community composition over the same range of resource supply parameters, when we initiate each pair of species in the system. To make both figures, all simulations were run until each state variable differed from one period to the next by no more than 10^−6^, or, if this threshold for stationarity was never reached, for 5,000 periods. Species were considered to be persisting if their population over the last period of the simulation was ever above 0.001. Species persistence was confirmed using invasion analysis. We see that, when the gleaner is absent from the system, the opportunist out-competes the supertunist over the entire parameter range. However, with the gleaner present, the opportunist and supertunist are able to persist over a relatively large parameter range. Note that, in the entire parameter range where the three species all coexist, the system displays non-simple community assembly, since the opportunist-supertunist two-species system collapses.

#### S2.3 Five species

Figure S4 shows community composition for the five species system shown in Fig 5 over a range of resource parameters. All simulations were run for 10, 000 time steps. A consumer was considered to be persisting if its population exceeded 0.001 at any point over the last cycle of the simulation. The panel on the left shows the precise species composition, while the panel on the right shows the number of surviving species. Note that the species with the lowest *R*^*^value (the gleaner, shown in orange in Fig 5) persists across all parameter values.

### S3 Three consumers with type II (Monod) per capita growth rates

#### S3.1 Simulations

We illustrate a system with three consumers, all with per capita growth rates given by type II (Monod) functions. Resource supply is continuous but fluctuating with *S*(*t*) = *A*(*α* sin(2*πt/P*)+ 1). The parameters are *r*_1_ = 0.85, *k*_1_ = 0.0085, *r*_2_ = 1.155, *k*_2_ = 0.05, *r*_3_ = 1.63, *k*_3_ = 0.244, *d*_*i*_ = 0.1 for all *i, q*_*i*_ = 1 for all *i, A* = 8, *α* = 0.9, *P* = 4.5, *w* = 0.5. The per capita growth rates are shown in fig S5. Simulations of the three-species system and each two-species system are shown in fig S6.

#### S3.2 Taylor expansions

The expansions of the per capita growth rates of each consumer are shown in fig S7. The expansions show that large deviations of resource abundance below the maximum are more detrimental to the gleaner, medium deviations are more detrimental to the opportunist, and small deviations are more detrimental to the supertunist. This is reflected in the resource distributions in fig S6. In each of the two-species systems, the resource distribution benefits the missing species, with the peak of the resource distribution occurring higher when the supertunist is invading, lower when the gleaner is invading, and at an intermediate value when the opportunist is invading.

### S4 An example with polynomial per capita growth rates

In the main text, we introduce a three-species system where each consumer has a polynomial per capita growth rate (Fig 2). We take

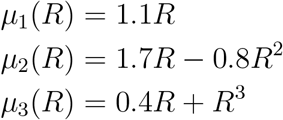

and *d*_*i*_ = 0.1 for all *i, q*_*i*_ = 2 for all *i*, and *w* = 0.5. Resource supply is given by *S*(*t*) = *A*(1.1 \**α* sin(2*πt/P*))^2^, with *A* = 1.5, *α* = 0.9, and *P* = 50. We choose this form for resource supply, rather than the sinusoidal form we have used previously, to provide higher skewness that facilitates the survival of the supertunist (see analysis in the main text).

For any *Z*, we can rewrite the per capita growth rates as polynomials centred around *Z*:

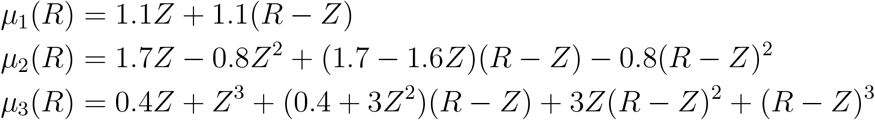

As in the main text, if we take 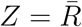 and take averages, we obtain

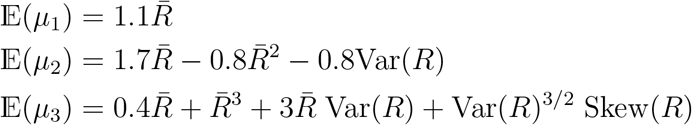

where Var(*R*) is the variance of the resource distribution and Skew 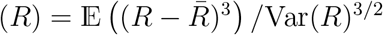 is its skewness. We use skewness here rather than the unstandardised moment 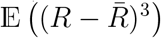 since its interpretation is more familiar, and since there are only three moments in the expression.

Using Condition 1, when all three species are coexisting together in the system, the system of equations above determines the mean, variance and skewness of the resource distribution: 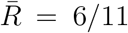, Var(*R*) = 0.1116, Skew(*R*) = 0.9919. This makes it possible to directly compare the contribution each moment makes to each species’ average growth rate at equilibrium (Fig 2B).

Condition 2 tells us when each species can invade the system from low numbers. The opportunist can invade if 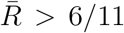, where the average is calculated with the opportunist absent from the system. The gleaner can invade if 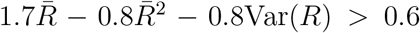. Thus, the gleaner will persist in the system if the mean resource is high enough (but not so high that the 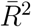 term takes over) and its variance is low enough. The supertunist can invade if 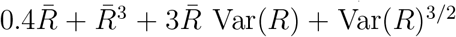. Using these conditions, we can investigate the assembly of the three-species system from each of the the smaller one-or-two-species systems.

We describe one pathway for community assembly (fig S9 shows simulations of the three-species system and each of the two-species subsystems). When there are no species in the system, the mean, variance and skewness of the resource distribution are as follows: 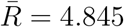, Var(*R*) = 17.183, Skew(*R*) = 0.3792. This allows the opportunist and the supertunist to invade, but not the gleaner, meaning the gleaner cannot survive alone in this system. When only the opportunist is in the system, from Condition 1, we know that 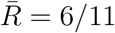. Substituting this into the invasion conditions for the gleaner and supertunist, we find that the gleaner can invade if Var(*R*) *<* 0.1116, and the supertunist can invade if Var(*R*)^3*/*2^ Skew(*R*) + 1.6364Var(*R*) *>* 0.2195. In this system, the resource variance is 0.1754 and skewness is 2.0214. This allows the supertunist but not the gleaner to invade. With the opportunist and the supertunist in the system, the variance falls to 0.1085. Since the opportunist is in the system, 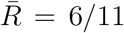, the invasion condition for the gleaner is the same as above. Thus, the gleaner can invade to reach the stable three-species system.

The intuition behind the differing requirements of the gleaner and supertunist from the moments of the resource distribution (as well as their differing impacts on the moments when they enter the system) can be readily seen from the species’ per capita growth rates (Fig 2A). The supertunist benefits from large deviations in resource abundance above the mean (captured by resource skewness) and the gleaner has negative growth when resources deviate too much on either side of the mean (captured by variance). Note that the supertunist’s average growth rate depends on all three of the average, variance and skewness of the resource distribution. As such, even when the opportunist is in the system setting the average resource level, the invasion of the supertunist depends on both the resource variance and skewness. Similarly, once the supertunist enters the system, the resource variance and skewness will change to balance the equation given by Condition 1. Thus, the supertunist cannot be said to consume a specific moment of the resource distribution. In models with realistic (i.e. non-polynomial) per capita growth rates, such as those we explore later in the main text, Conditions 1 and 2 will usually exhibit many degrees of freedom, making it impossible to summarise a consumer’s niche exactly in terms of one or two moments of the resource distribution. In this way, realistic systems exhibit far more nuance than the toy case explored by Levins *(23)*.

In fig S8, we show the expressions obtained when we take *Z* = *R*_max_ and *Z* = 0. Note that taking *Z* = 0 gives a particularly straightforward characterisation of the niches of the three species in terms of moments, since, in this case, the term corresponding to the second moment vanishes in the expression for the supertunist. This illustrates the utility that can be gained by centering the expression for the average per capita growth rate around a particular value of *R*.

Explicitly, when we take *Z* = 0, we get the following:

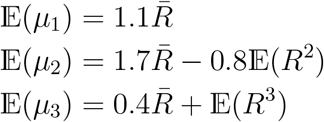

If the opportunist is in the system, so that 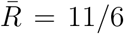, Condition 2 tells us that the gleaner can invade if 𝔼 (*R*^2^) *<* 9*/*22 and the supertunist can invade if 𝔼 (*R*^3^) *>* 21*/*55. Thus, while the opportunist consumes the average resource abundance 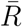, the gleaner is harmed by 𝔼 (*R*^2^), and the supertunist consumes 𝔼 (*R*^3^).

### S5 Deriving the Taylor expansions for type II (Monod) and type III per capita growth functions

To derive the Taylor expansions, we will use that fact that

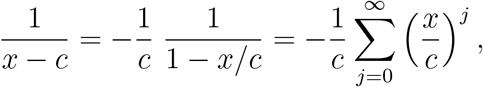

which converges for |*x*| *<* |*c*|.

We can apply this directly to the Monod function *μ*(*R*) = *rR/*(*k* + *R*):

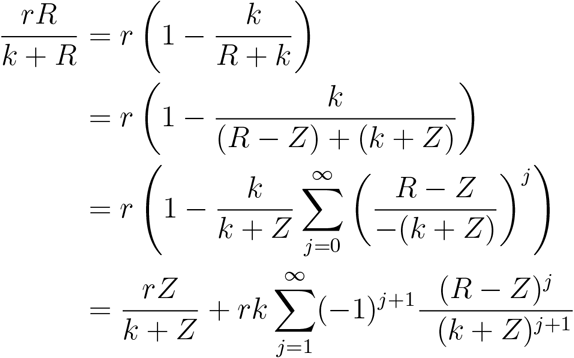

This converges for | *R* − *Z*| *< k* + *Z*. Since resources are always positive, this condition is vacuous for *R < Z*. Thus, the Taylor expansion converges for *R < k* + 2*Z*.

To derive the Taylor expansion for the type III function *μ*(*R*) = *rR*^2^*/*(*k* + *R*^2^), we proceed similarly:

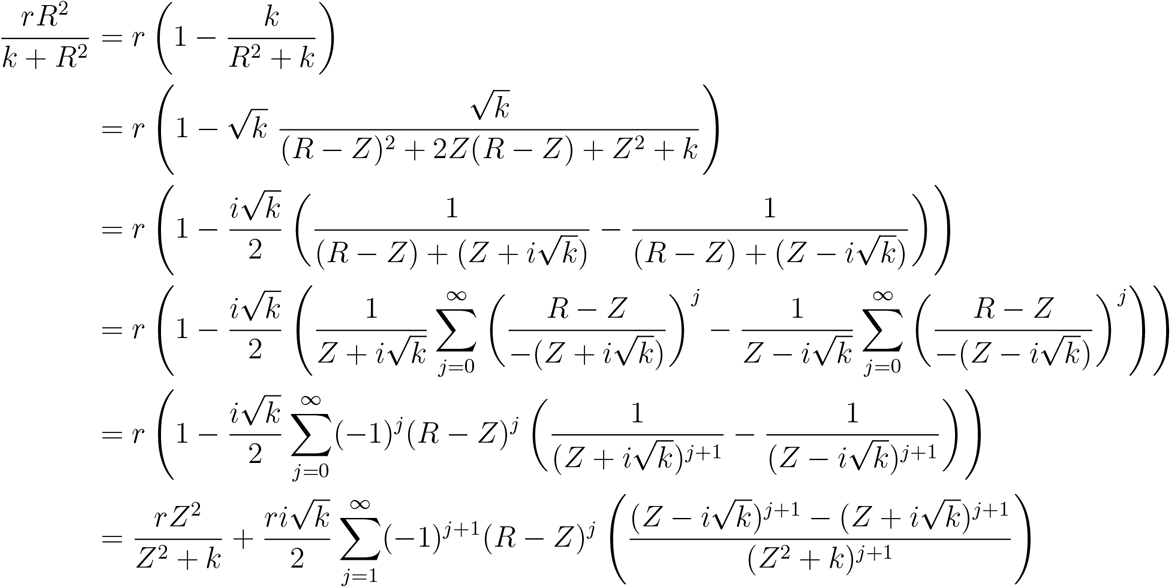

This converges for 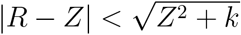. As with the Monod function, this condition is vacuous for *R < Z*, so the condition amounts to 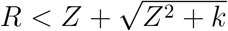.

### S6 Expressing the average per capita growth rates in terms of moments

Suppose *μ*(*R*) is analytic around the point *R* = *Z*, with Taylor expansion given by

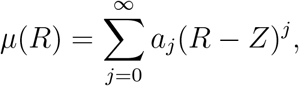

and radius of convergence *r*. We claim that, as long as we can find *ρ < r* such that, for all *t >* 0, we have |*R*(*t*) − *Z*| ≤ *ρ*, we can calculate the expected value of *μ* term-by-term:

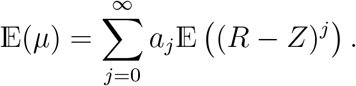

To see this, recall that 𝔼 (*μ*) is given by

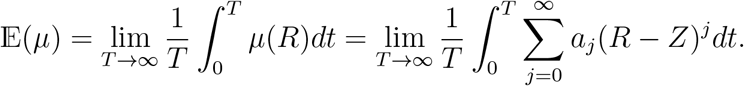

We will begin by showing that we can take the integral inside the sum. Given *ρ* satisfying the hypotheses above, choose 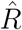 such that 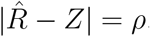. For any *T >* 0 we have:

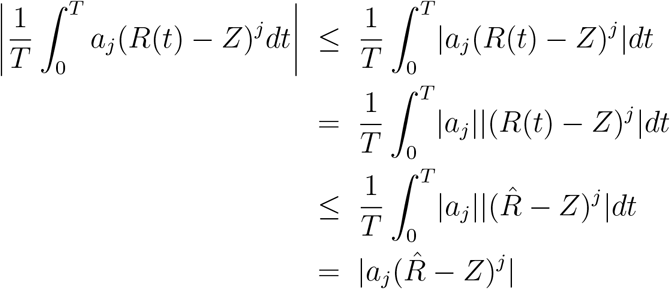

The first inequality is the triangle inequality and the second follows from our assumption that |*R*(*t*)\**Z*| − *ρ*. Now, write 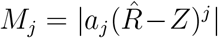. Since 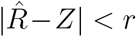, we know that 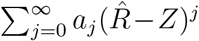 converges absolutely, meaning ^−^ converges. Thus, the sum of the integrals converges and we have

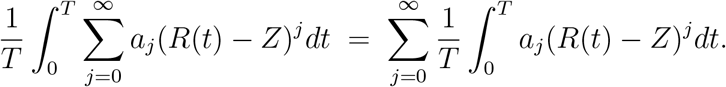

Now, we are assuming that the limit defining 𝔼 ((*R* \**Z*)^*j*^) exists for all *j*. Thus, by Tannery’s theorem *(58)*, to establish the equality

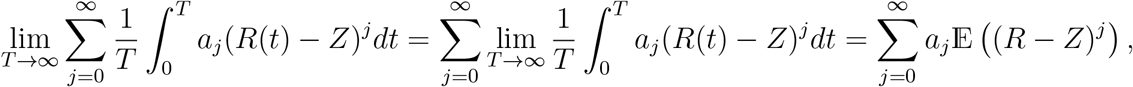

we need only bound the summand on the left-hand side of the equation. For this, we can take *M*_*j*_.

### S7 An example that satisfies the small variance approximation

To illustrate the limitations of the small variance approximation, we describe its use in a classical gleaner-opportunist system. The opportunist has a linear per capita growth rate *μ*(*R*) = 0.358*R*, and the gleaner has Monod per capita growth rate *μ*(*R*) = *rR/*(*k* + *R*) with *r* = 8 and *k* = 20 (fig S10A). Resource supply is given by *S*(*t*) = *A*(*α* sin(2*πt/P*) + 1) with *A* = 1.5, *α* = 0.9, *P* = 11 and *w* = 0.5, and both species have *d*_*i*_ = 0.1 and *q*_*i*_ = 1.

Since the opportunist has a linear per capita growth rate, it consumes the mean of the resource distribution *sensu* Levins *(23)* (cf. species 1 in Fig 2A and the corresponding description in the text). Thus, if the opportunist is in the system, it determines the mean of the resource distribution 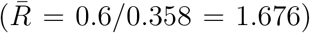. If we assume the small variance approximation holds, then the gleaner’s average per capita growth rate is described by the second order truncation of Equation 6:

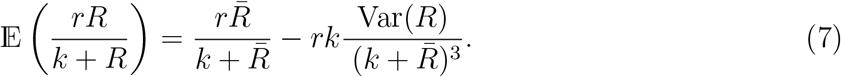

Note that, with the opportunist in the system setting the resource mean, the only degree of freedom in this equation is the resource variance. Thus, by Condition 2, the gleaner can only invade the opportunist monoculture if the variance in the resource distribution is sufficiently low (cf. the gleaner in Fig 2A). In this way, the system reduces to the simple polynomial case described by Levins *(23)* and recalled in the preceding section.

Equation 6 for the gleaner in this system is as follows:

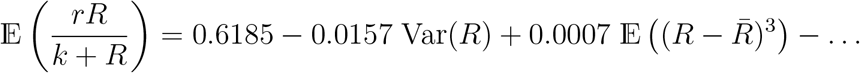

The central moments in this system increase non-monotonically; however, the coefficients in the equation above decrease much more rapidly, and so we may neglect all but the first two terms in this equation (see fig S10E), giving us

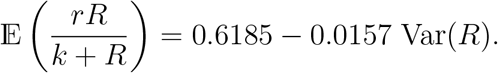

Thus, the gleaner can invade as long as the variance in the opportunist monoculture is less than 1.178. This is the case, meaning the gleaner is able to invade. Note, however, that once the gleaner is in the system, the approximation above tells us the variance should be 1.178. In practice, the variance in the two-species system is 1.194, showing that, even in this case, the small variance approximation loses relevant information.

### S8 Exploring the graphical representations of the average per capita growth rate expansions

In this section, we elaborate on the graphical representations of the expansions of the average per capita growth rates for the gleaner-opportunist system (Figs 3D and 3E) and the gleaner-opportunist-supertunist system (Fig 4) from the main text. Figs S11 and S13 compare the expansions calculated in the full gleaner-opportunist system to the expansions calculated in each monoculture; for convenience, we reproduce Figs 3D and 3E here. The similar shape of the plots in each case indicates that the specialisation the consumers display on different moments of the resource distribution is consistent whether they are invading or resident. Note that, when a species is resident, its average per capita growth rate is equal to *d*_*i*_ + *w* by Condition 1, which in this system is 0.6. However, when a species is invading its per capita growth rate is greater than 0.6. This is reflected in panels A and C of Figs S11 and S13, although the scale on the axis makes the difference hard to read off the plot. A similar plot for the gleaner-opportunist-supertunist system is shown in fig S16.

To illustrate the cumulative contribution of terms in the average per capita growth rate expansions, figs S12 and S14 show the remainder after the expansion is calculated to *n* terms, starting with *n* = 0 when the remainder is equal to the whole of the average per capita growth rate. fig S12B shows clearly that the combined sum of the third and higher terms is more negative for the gleaner than the opportunist; in fact, as noted in the main text, this is also true for the sum of terms starting at the second, although the scale of the plot makes this difficult to read off. In this way, higher moments in the resource distribution are more harmful to the gleaner than the opportunist. The analogous plot for the gleaner-opportunist-supertunist system is shown in fig S17.

Finally, Figs S15 and S18 show plots of the moments in these systems alongside the coefficients of the average per capita growth rate expansions. Note that the moments that appear in the expansions are not normalised: if | *R*− *Z*| is always less than 1, as in the gleaner-opportunist-supertunist system where resources fluctuate strictly between 0 and *K* = 0.5, then the higher moments will vanish, while if, as in the gleaner-opportunist system, |*R* − *Z*| is often greater than 1, the higher moments will rapidly increase. This is balanced by the coefficients in the expansion, which rapidly decrease in Figs S15B and S15D. Note that in the gleaner-opportunist-supertunist system, where we have chosen to expand around *R* = *K*, the coefficients in the expansions are independent of the resource distribution in the system.

### S9 A gleaner-opportunist system with discrete resource supply

We give an example of a gleaner-opportunist system where resources are refreshed at discrete time intervals (e.g. in an experimental serial transfer regime) *(9, 59)*. Between pulses, the system evolves as usual, except with *w* = 0 and no resource supply:

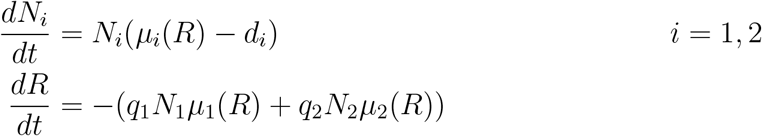

At intervals of length *T*, the resource is instantaneously supplied at a constant amount *S*_0_. In this system, we take *T* = 10 and *S*_0_ = 0.6. The consumers both have Monod per capita growth rates with parameters *r*_1_ = 1, *k*_1_ = 0.5, *r*_2_ = 2.5, *k*_2_ = 1.7, *d*_*i*_ = 0.1 for all *i*, and *q*_*i*_ = 2 for all *i*. The per capita growth rates are shown in fig S19A.

In both the two-species system and the monocultures, resources are fully depleted between pulses. Thus, the maximum resource abundance in each system is *R*_max_ = *S*_0_ = 0.6. Since *R*_max_ is fixed, we take Taylor expansions around *Z* = *R*_max_ as well as 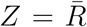. The resulting expressions for the average per capita growth rates in the two-species system are shown in Figs S19D and S19E. From these, we can see that the opportunist is advantaged by higher moments in resource supply relative to the gleaner. We can form the same plots in either monoculture, but we have not shown them here since they are indistinguishable from Figs S19D and S19E; this reflects the fact that, whether one species is invading and the other is resident, or both are resident, the species in this system partition the moments of the resource distribution in the same way. Thus, the two species coexist robustly, the gleaner benefiting from the lower moments in the resource distribution and the opportunist from the higher moments.

### S10 The gleaner-opportunist-survivalist system

Here we consider the system obtained by replacing the supertunist in the system illustrated in Fig 1 by a consumer whose per capita growth rate has the same type III functional form as the supertunist, but with a lower maximum growth rate and a lower death rate (fig S20A). We refer to this consumer as a survivalist. As described in(37), one biological mechanism that can give rise to a per capita growth rate of this shape is a dormancy strategy, whereby the consumer enters a dormant state with a low death rate when resources are scarce, switching to an active state when resources are abundant. Fig S21 is the analogue of Fig 1 for this system, showing simulation results for each of the two-species systems. The survivalist is a poor competitor when resources are high, and the superior competitor when resources are low (fig S20A). However, since its growth rate is still negative in the resource range where it is effectively the superior competitor, the survivalist relies on periods of high resource abundance that allow it to achieve a positive growth rate, alongside periods of low resource abundance that reduce the populations of its competitors. Thus, as is the case for the supertunist, the survivalist benefits from peaks in resource availability coupled with periods of low resource availability. This overlap in the niche of the survivalist and the supertunist can be seen from the similarity between the expansion of the survivalist’s average per capita growth rate in fig S20B and the supertunist’s in Fig 4.

### S11 Community assembly

In most of the examples we have investigated, not all subsets of species are able to coexist. Often, one species acts as a keystone species that facilitates the coexistence of all of the others. For example, in the five-species system illustrated in Fig 5, the species with the lowest *R*^*^value (the gleaner, shown in orange in Fig 5) assumes the role of the keystone species. This reflects the species’ feedback on the resource distribution; when it is in the system, it increases resource fluctuations, creating niches for the other species. When the orange species is absent, the system collapses and only the species with the second lowest *R*^*^ value (the light blue species in Fig 5) survives.

In the invasion graph below, we illustrate community assembly in the five-species system. The species are numbered 1 to 5 in terms of increasing *R*^*^ value, so that the orange species in Fig 5 is species 1 and the dark blue species is species 5. Communities that coexist are coloured green and those that collapse are red. The arrows indicate which species can invade each community. If, in the five-species community, a single species is reduced to low numbers, the arrows indicate how the community recovers (cf. the invasion graphs studied by Hofbauer and Schreiber *(60)* and the assembly graphs studied by Song *(61)*). When species 3, 4 or 5 are at low numbers, the remaining four species coexist (second row in the invasion graph), and the missing species is able to invade from the system’s new stable orbit. When species 2 is at low numbers, species 4 also declines to low numbers and the system approaches the stable orbit of the community consisting of species 1, 3 and 5. From this state, species 2 can return, followed by species 4. When the keystone species 1 is reduced to low numbers, the community approaches the stable orbit of the species 2 monoculture. From there, species 1 can reinvade, followed by the remaining species.

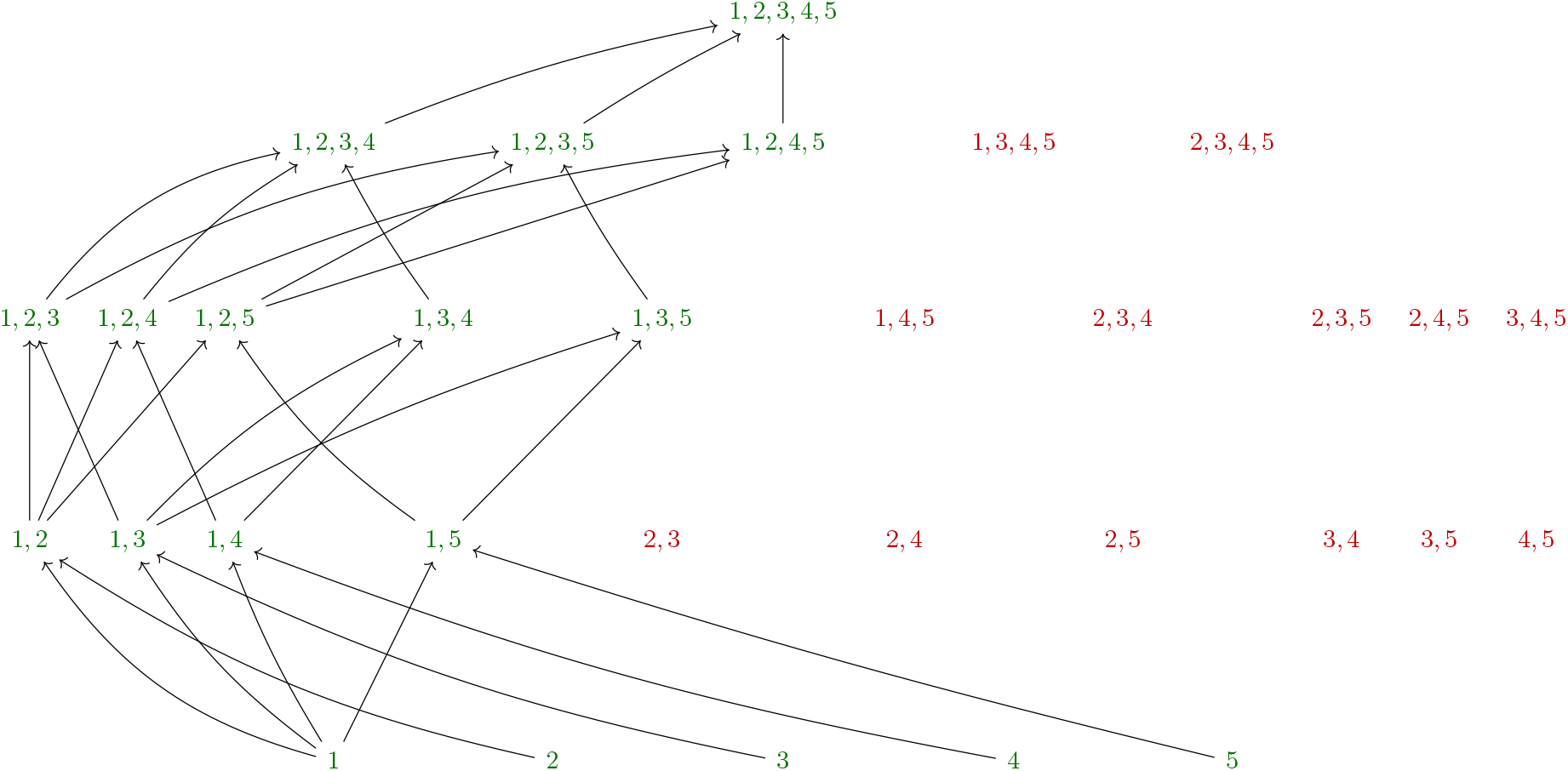

Note that we have used invasion tests to construct the graph above. If any of the communities display alternate stable states then the analysis will not be valid; however, we have also explored multiple different initial values with simulations and found no evidence of multiple stable states.

The graph below represents the competitive outcomes for each two-species community. If species *i* excludes species *j* we denote this by a red arrow *i* → *j*, and if species *i* and *j* coexist we denote this by a green arrow *i* ↔ *j*.

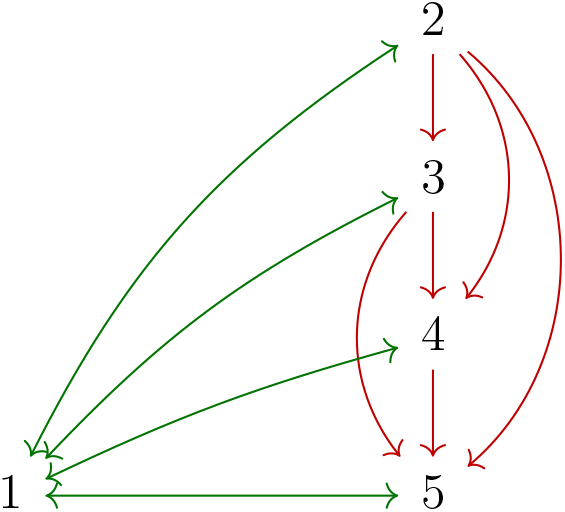

Consumers 2 *−* 5 display a strict competitive hierarchy. This is a feature this system has in common with other examples of emergent coexistence *(39)*. Note that this hierarchy is reflected in the invasion graph; in any sub-community, species higher up the competitive hierarchy can invade, until a cut-off is reached below which no species can invade.

As in the five-species system, in many of the systems we have investigated, the species with the lowest *R*^*^ value (the gleaner) is the keystone species. Note, however, that this need not be the case. We give two example systems, both consisting of consumers with per capita growth rates given by the following: *r*_1_ = 1.1, *k*_1_ = 0.54, *l*_1_ = 1, *r*_2_ = 1.85, *k*_2_ = 1.75, *l*_2_ = 1, *r*_3_ = 3.38, *k*_3_ = 7, *l*_3_ = 2. These are shown in fig S22.

In the first system, the resource is supplied, as in *S9 A gleaner-opportunist system with discrete resource supply*, in discrete pulses (*S*_0_ = 2.8) at intervals of *T* = 8. The death rates and quotas of the consumers are *d*_*i*_ = 0.3 for all *i*, and *q*_*i*_ = 2 for all *i*. Simulations of the three-species system and each two-species system are shown in fig S23, from which we can see that the supertunist in this system is the keystone species. In this case, where the supplied resource distribution is highly skewed, the supertunist pulls down the resource level and facilitates the survival of the gleaner.

Finally, fig S24 shows simulations for the system with the same per capita growth rates and quotas, but *d*_*i*_ = 0.085 for all *i*, and continuous resource supply *S*(*t*) = *A*(*α* sin(2*πt/P*) + 1), with *A* = 1.5, *α* = 1, and *P* = 30 and *w* = 0.5. In this system, all two-species systems coexist.

**Figure S1:**
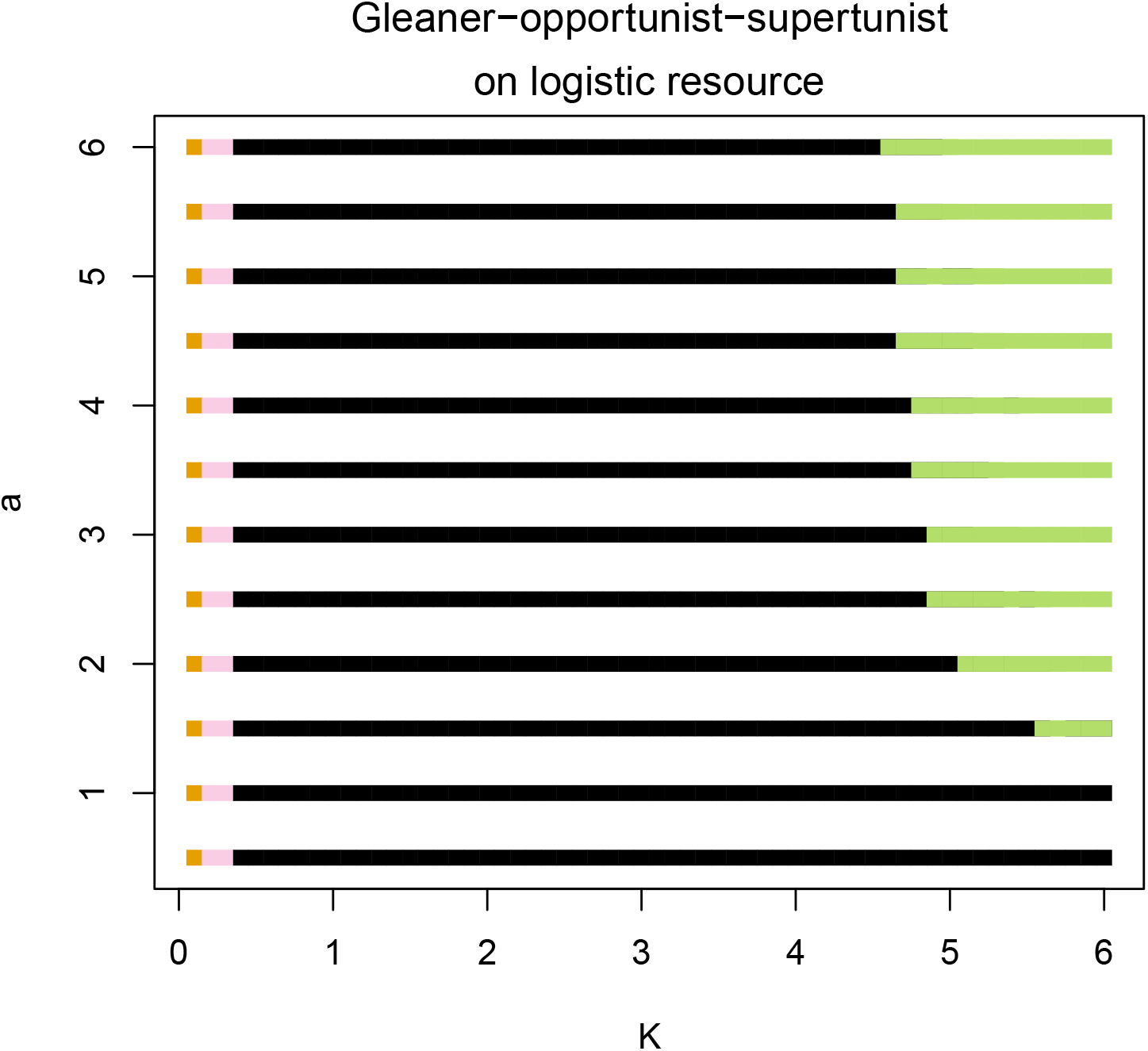
Structural stability for the gleaner-opportunist-supertunist system with logistic resource supply studied in the main text. Other than *a* and *K*, all parameters are fixed at the values given in *S1 Full models and parameter values from the main text*. The coexistence outcomes are as follows: gleaner monoculture is shown in orange; gleaner and opportunist in pink; opportunist and supertunist in green; three-species coexistence in black.

**Figure S2:**
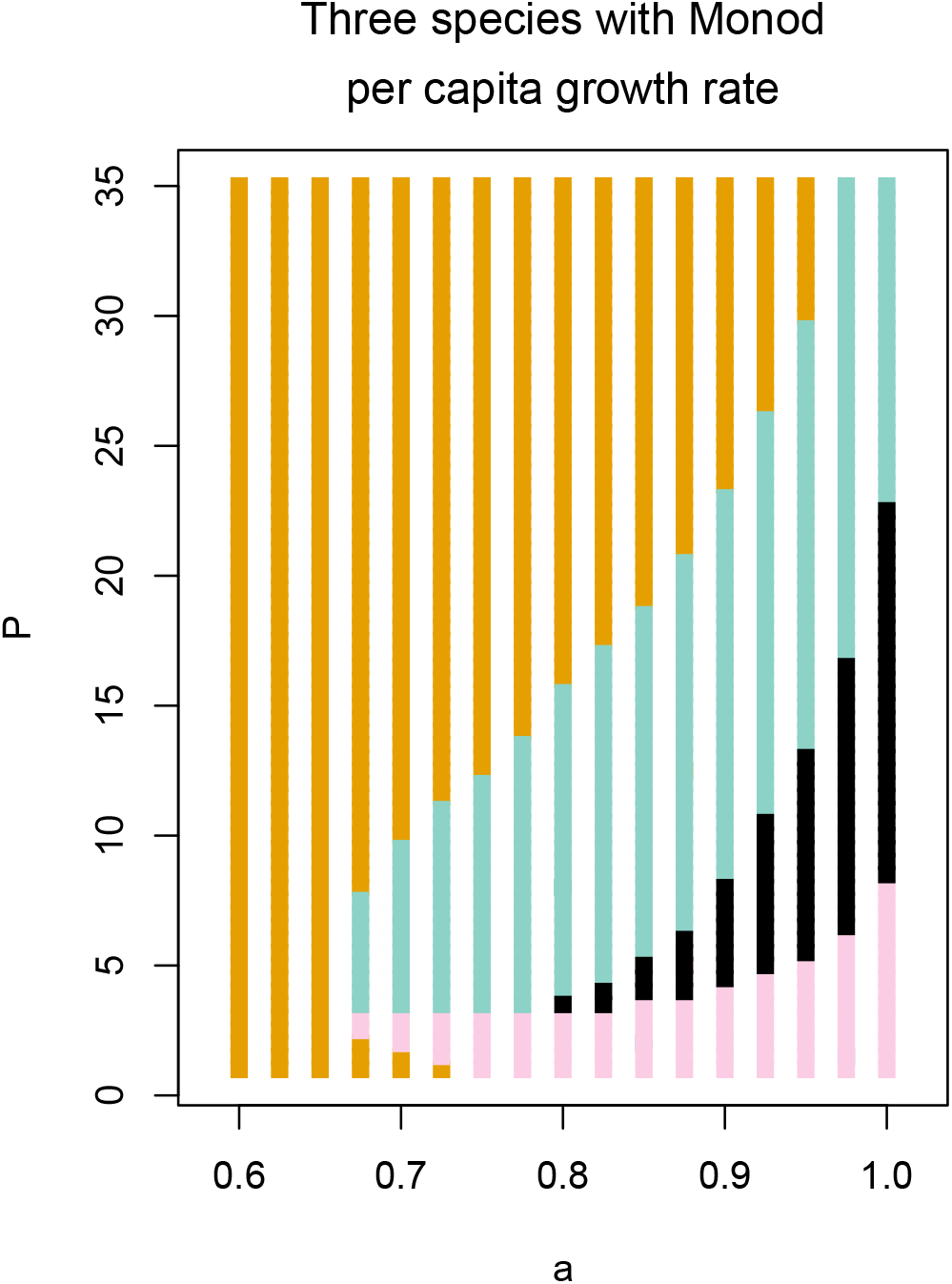
Structural stability for the gleaner-opportunist-supertunist system described in *S3 Three consumers with type II (Monod) per capita growth rates*. Other than *P* and *a*, all parameters are fixed at the values given in *S3*. The coexistence outcomes are as follows: gleaner monoculture (orange); gleaner and opportunist (pink); gleaner and supertunist (teal); three-species coexistence (black). Note the gleaner always persists.

**Figure S3:**
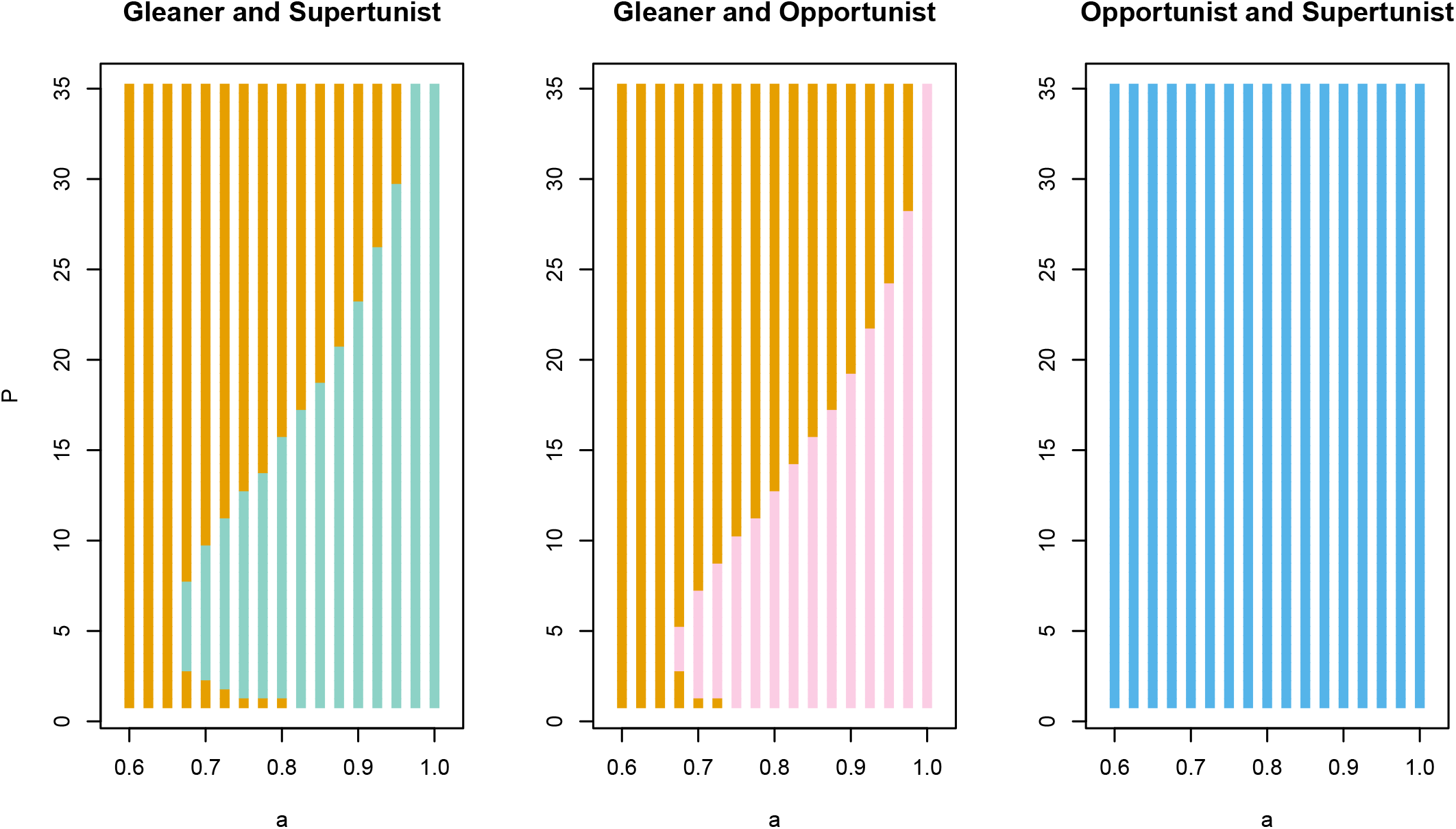
Structural stability for each two species subsystem of the gleaner-opportunist-supertunist system described in *S3 Three consumers with type II (Monod) per capita growth rates*. As in fig S2, gleaner monoculture is shown in orange; gleaner and opportunist in pink; gleaner and supertunist in teal. Opportunist monoculture is shown in blue.

**Figure S4:**
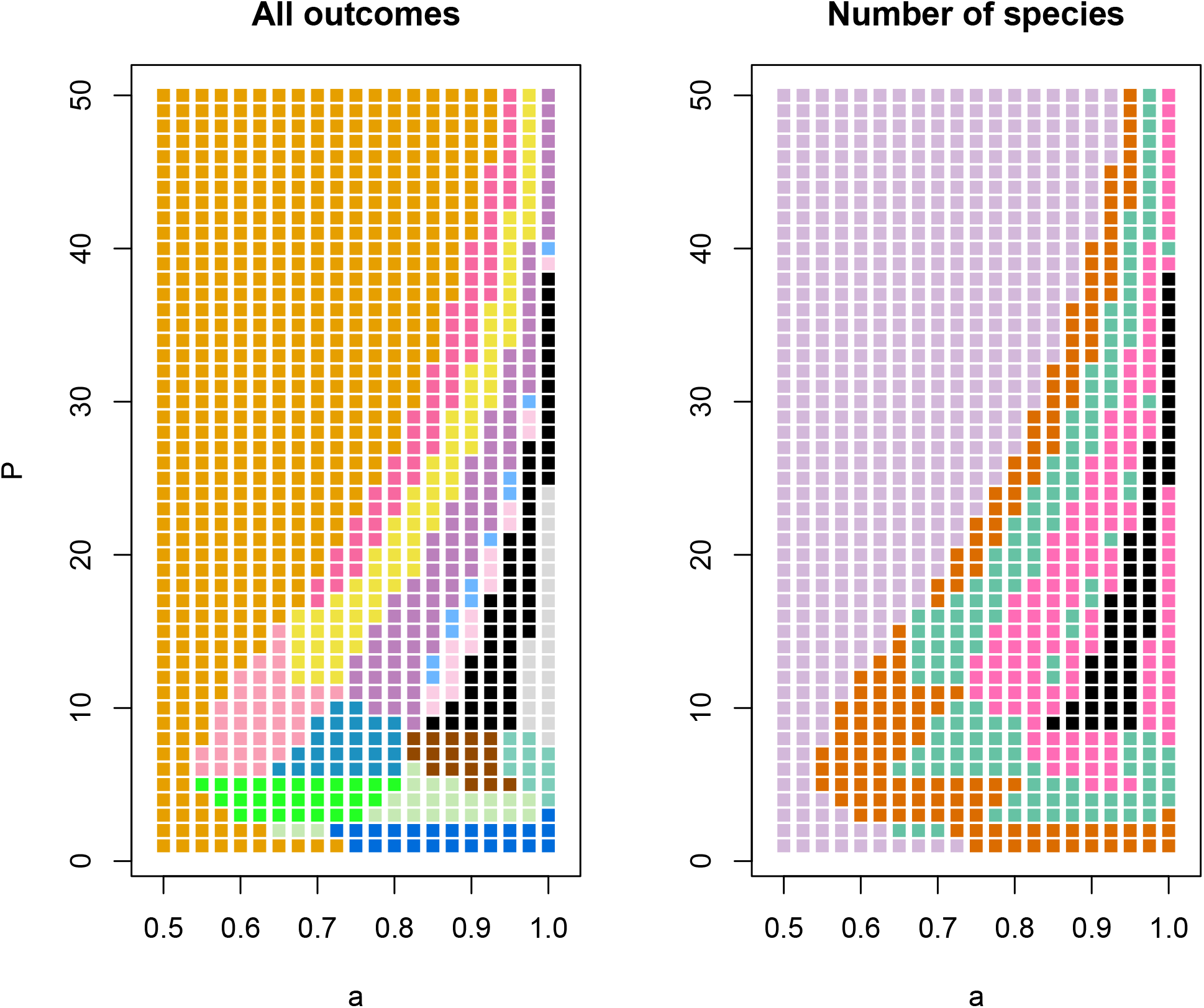
Structural stability for the five species system described in the main text. The left panel shows all coexistence outcomes, with five-species coexistence shown in black and gleaner monoculture shown in orange. The other colours are as follows, where, as in *S11 Community assembly*, we number the species by increasing *R*^*^ value: species 1 and 2; species 1 and 3; species 1 and 4; species 1 and 5; species 1, 2 and 3; species 1, 2 and 4; species 1, 3 and 4; species 1, 3 and 5; species 1, 4 and 5; species 1, 2, 3 and 4; species 1, 2, 3 and 5; species 1, 2, 4 and 5; species 1, 3, 4 and 5. Note that not all of the possible sub-communities occur. The right panel shows how many species survive in each of the outcomes: 1 species, 2 species, 3 species, 4 species and black for all 5 species.

**Figure S5:**
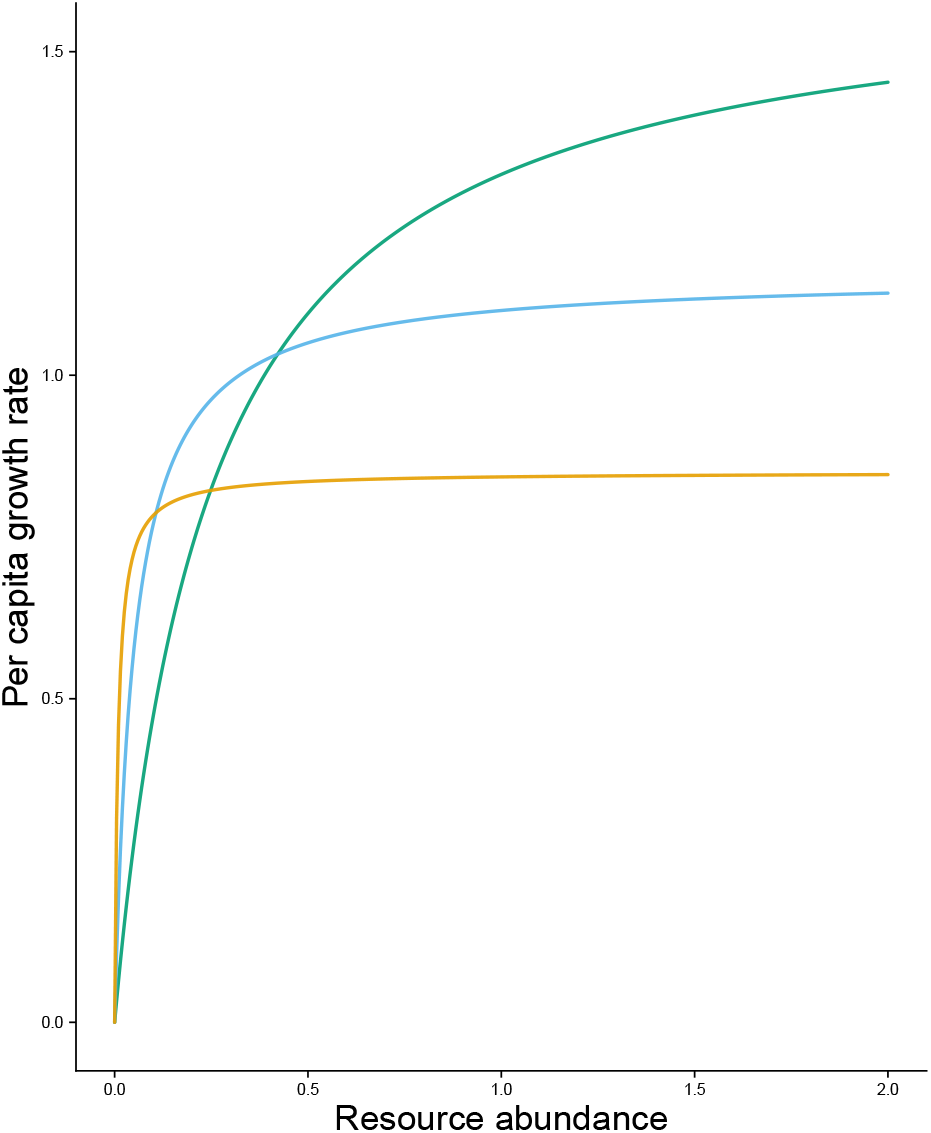
Per capita growth rates in the system described in *S3 Three consumers with type II (Monod) per capita growth rates*. As in previous examples, we refer to the orange consumer as the gleaner, the blue as the opportunist, and the green as the supertunist.

**Figure S6:**
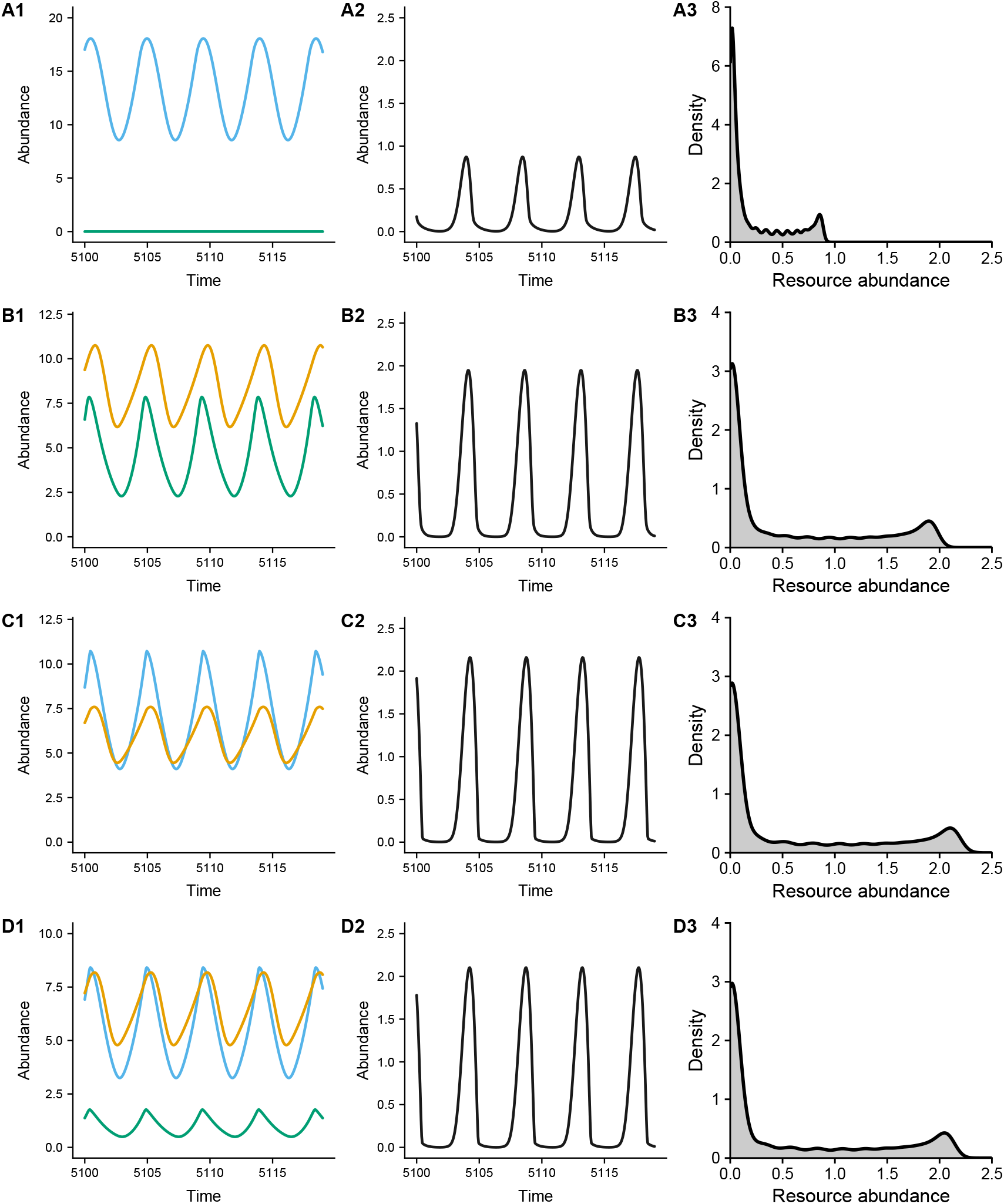
Simulation results for the system described in *S3 Three consumers with type II (Monod) per capita growth rates*. The opportunist is shown in blue, the gleaner in orange, and the supertunist in green. **A**. The opportunist-supertunist system. **B**. The gleaner-supertunist system. **C**. The gleaner-opportunist system. **D**. The complete three-species system.

**Figure S7:**
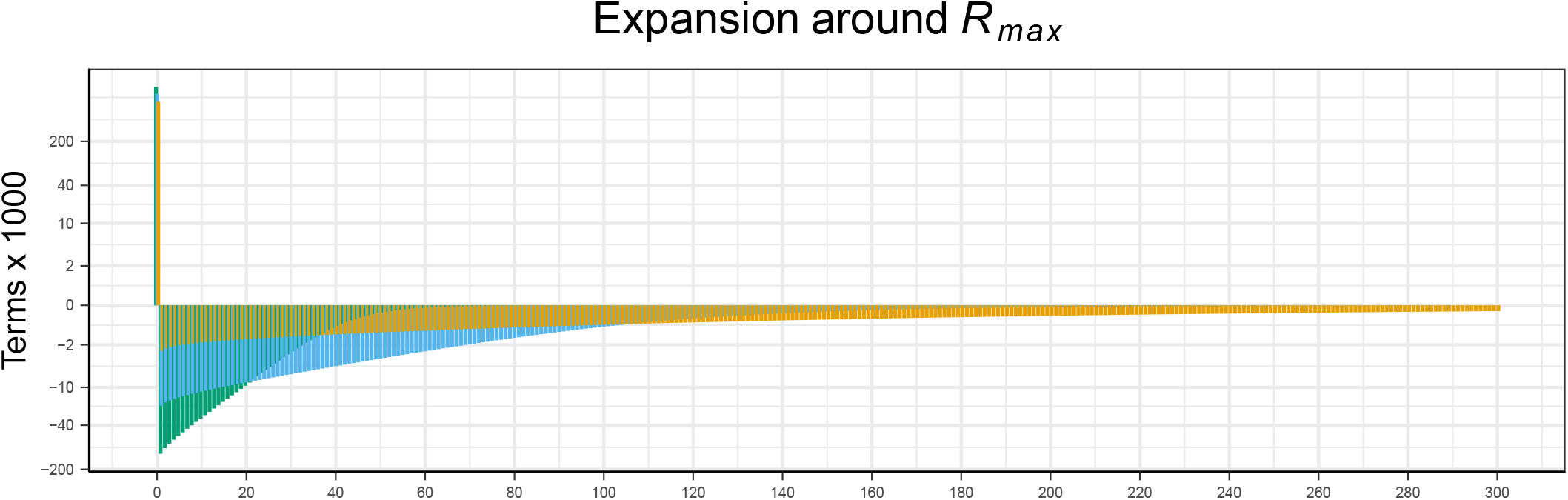
Expansions around *R*_max_ for the system described in *S3 Three consumers with type II (Monod) per capita growth rates*. Terms corresponding to the opportunist are shown in blue, those corresponding to the gleaner are in orange, and those corresponding to the supertunist are green.

**Figure S8:**
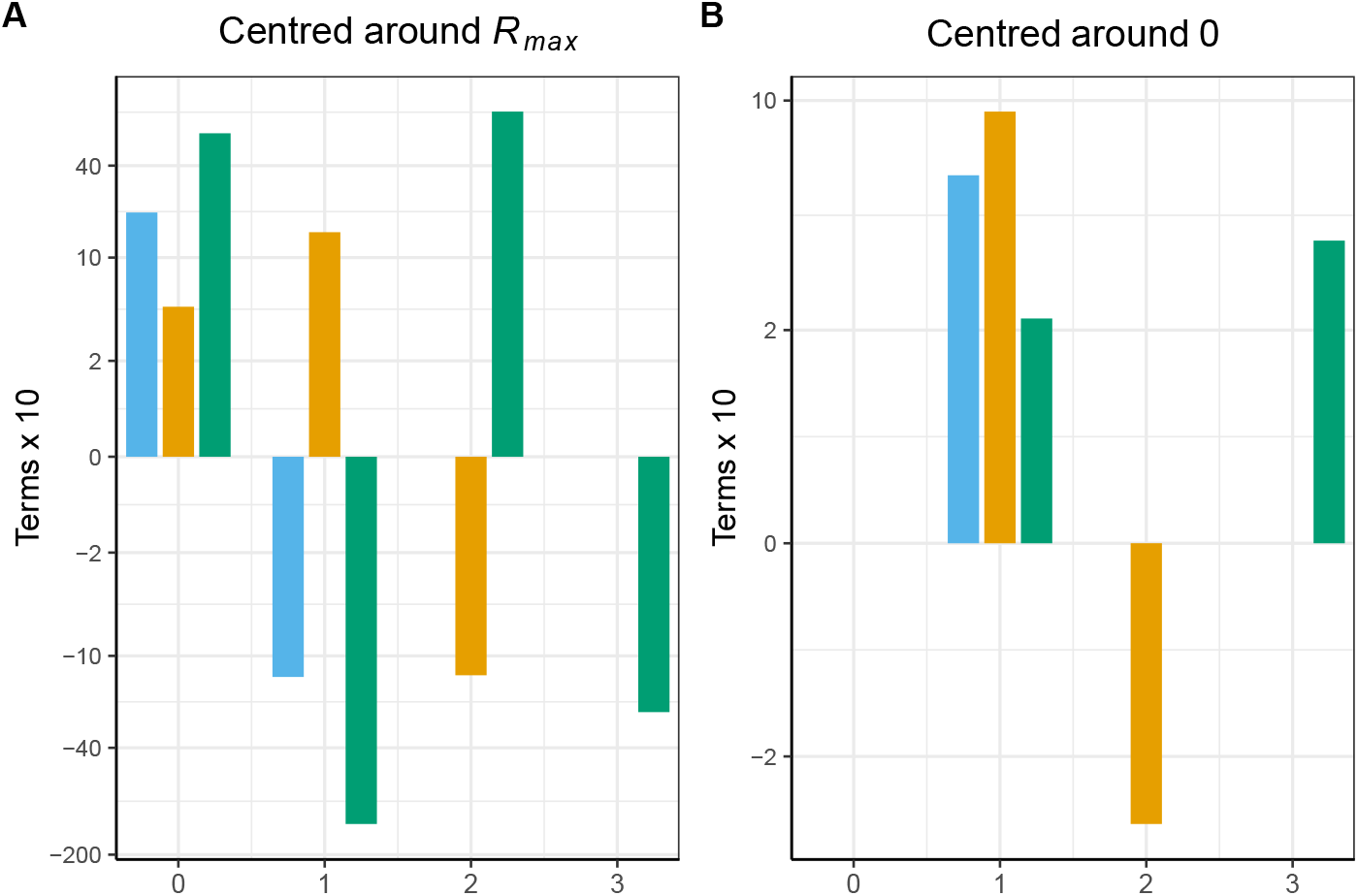
Terms in the average per capita growth rate expansions for the polynomial system shown in Fig 2; opportunist (blue), gleaner (orange), supertunist (green).

**Figure S9:**
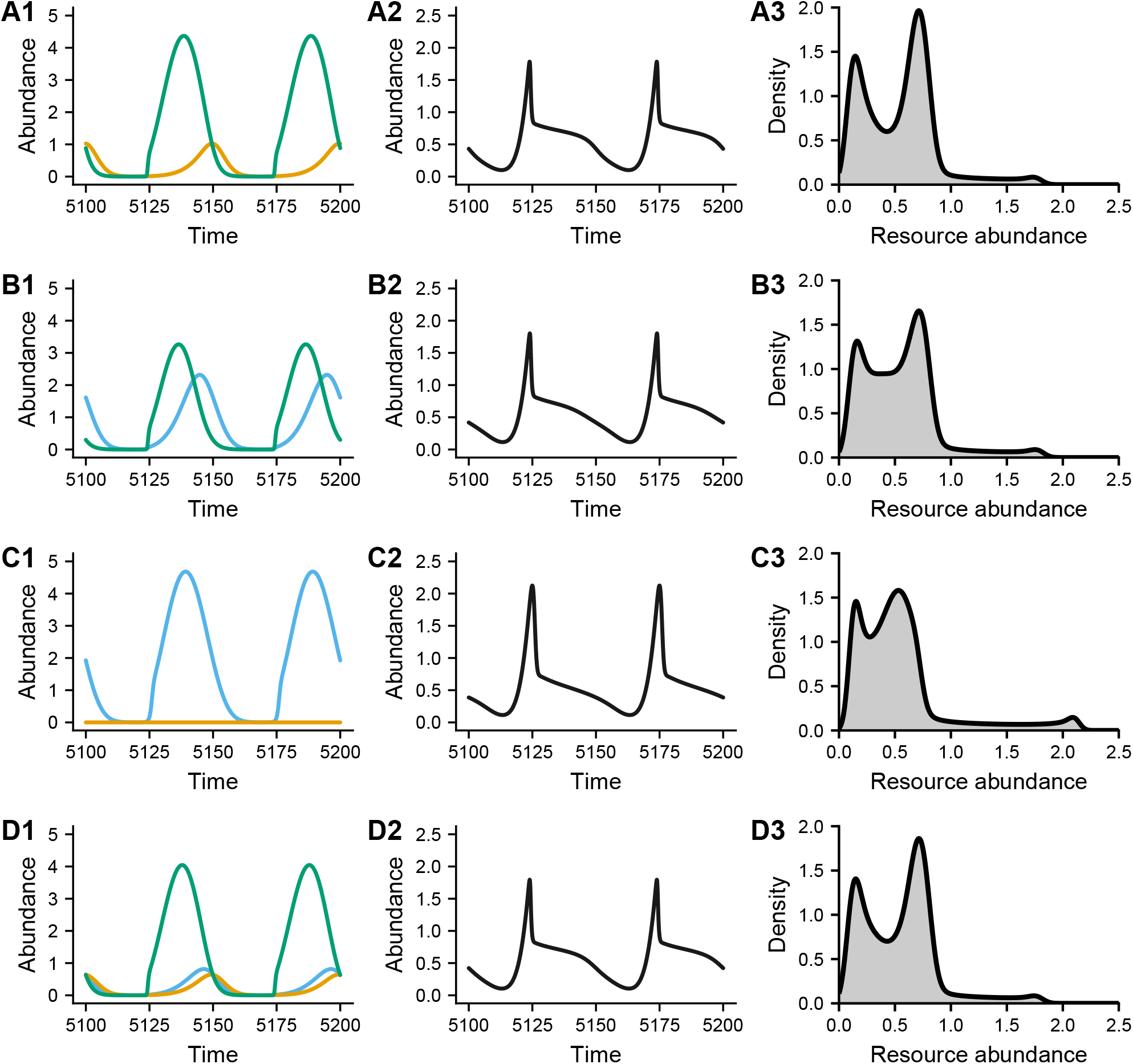
Simulation results for the polynomial system shown in Fig 2. **A**. The system consisting of only the gleaner and the supertunist. **B**. The system consisting of only the opportunist and the supertunist. **C**. The system consisting of only the opportunist and the gleaner. **D**. The complete three-species system.

**Figure S10:**
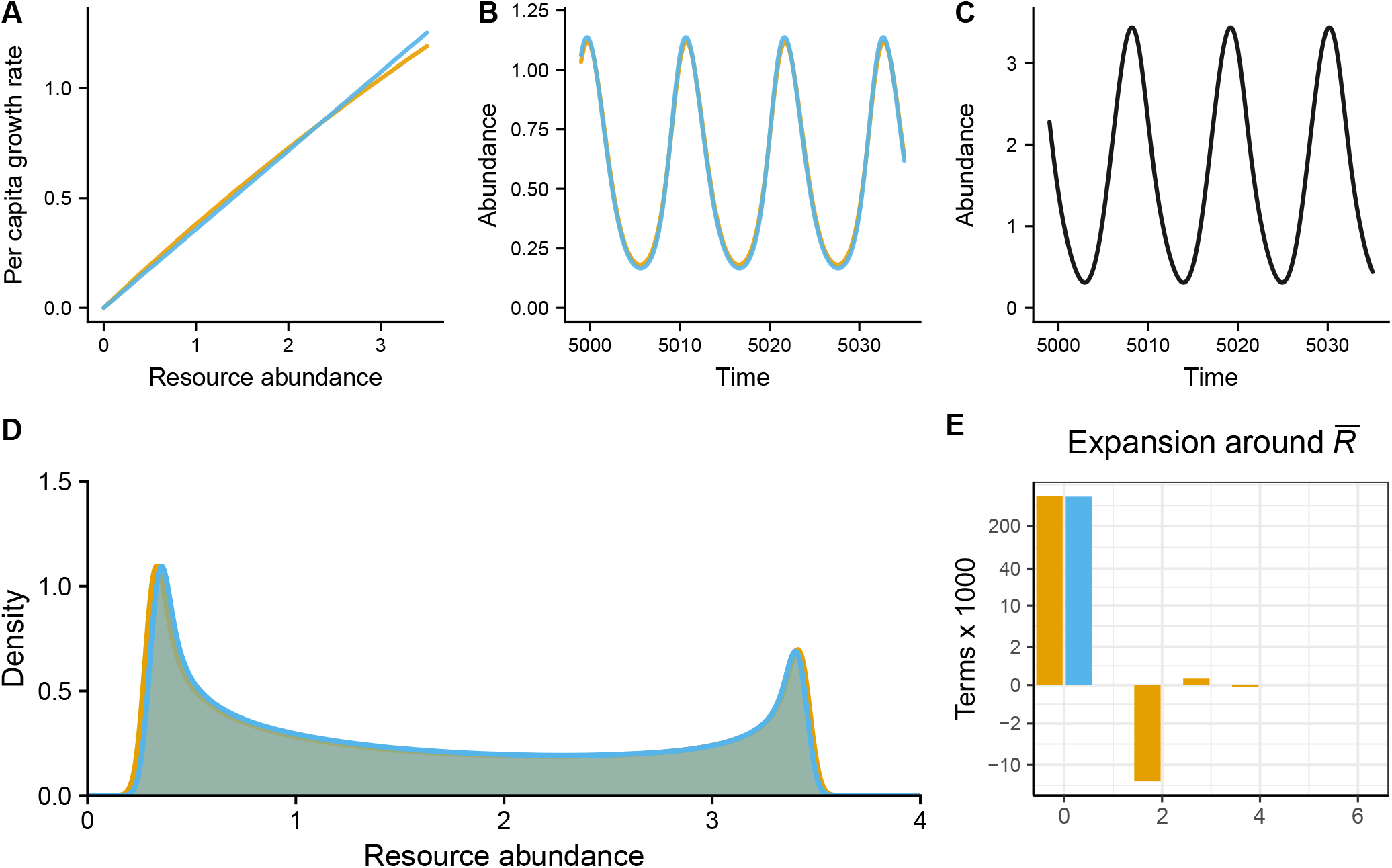
Simulation results for the gleaner-opportunist system in *S7 An example that satisfies the small variance approximation*. The opportunist is shown in blue and the gleaner in orange. Per capita growth rates. **B**. Simulation results showing steady-state population dynamics. **C**. Simulation results showing the corresponding resource dynamics. **D**. Kernel density estimates for resource abundance through time. The orange is in the gleaner monoculture system, the blue in the opportunist monoculture system. **E**. The terms in the expansions of each consumer’s per capita growth rate around 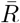 in the two-species system. Note the similarity between the two consumers, which is necessary to allow for a system that satisfies the small variance approximation (when at least one consumer has a type II per capita growth rate). In the analysis, we ignore all but the first two terms in the expression for the gleaner’s average per capita growth rate.

**Figure S11:**
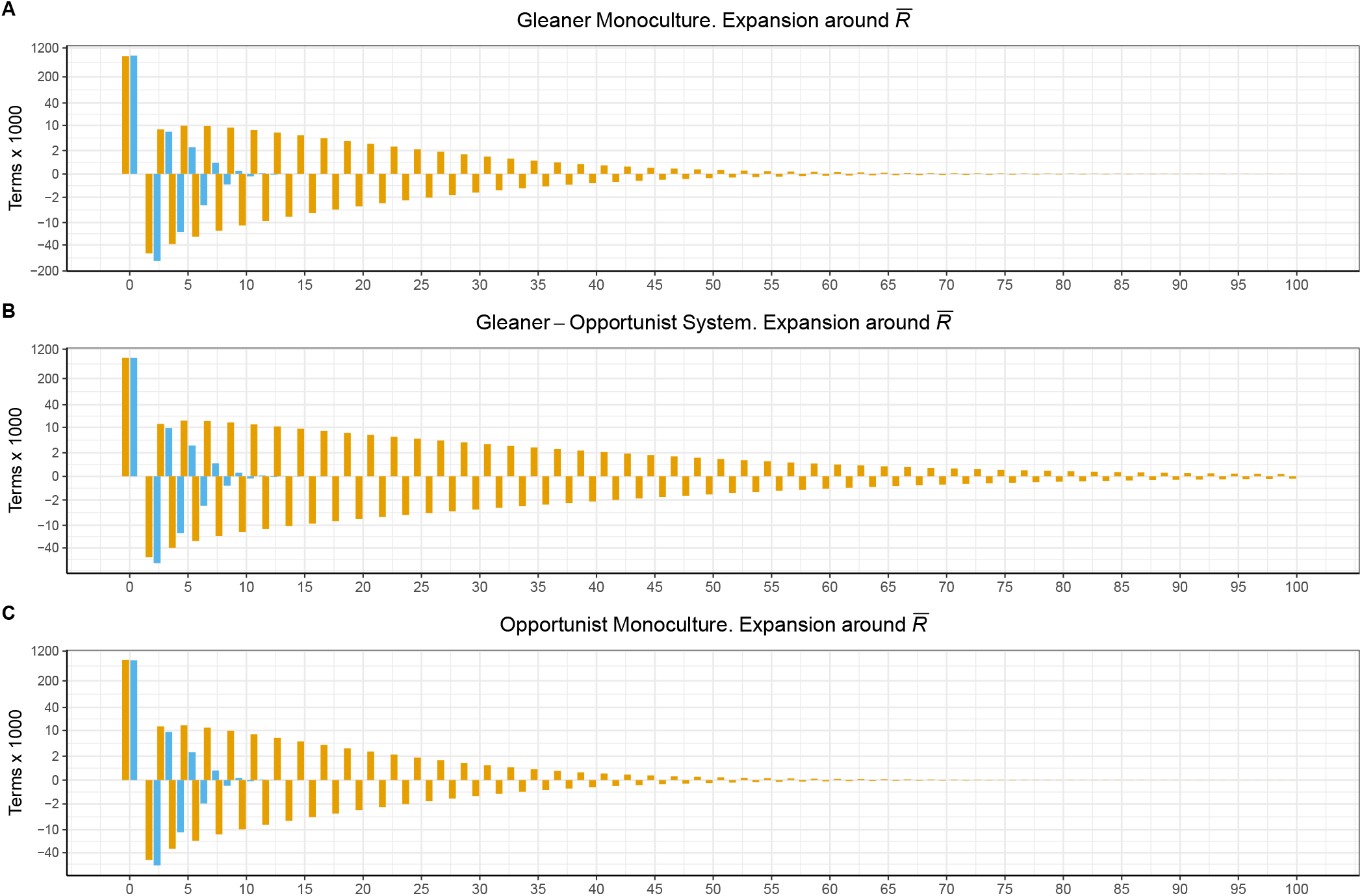
Expansions around 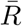 in the gleaner-opportunist system described in the main text. Terms corresponding to the opportunist are shown in blue and those corresponding to the gleaner are shown in orange.

**Figure S12:**
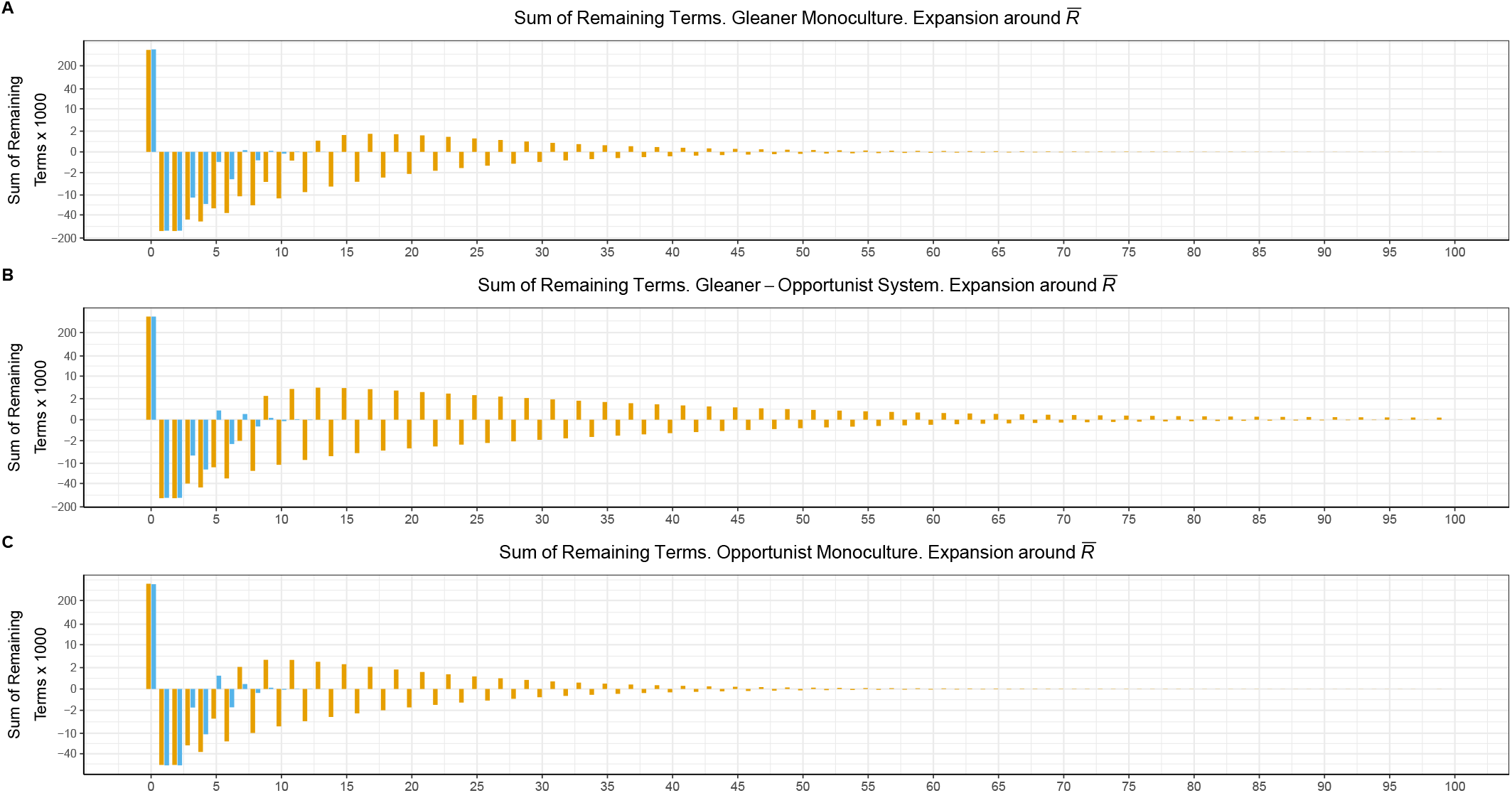
Remainders in the expansions around 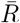 in the gleaner-opportunist system described in the main text. Terms corresponding to the opportunist are shown in blue and corresponding to the gleaner are shown in orange.

**Figure S13:**
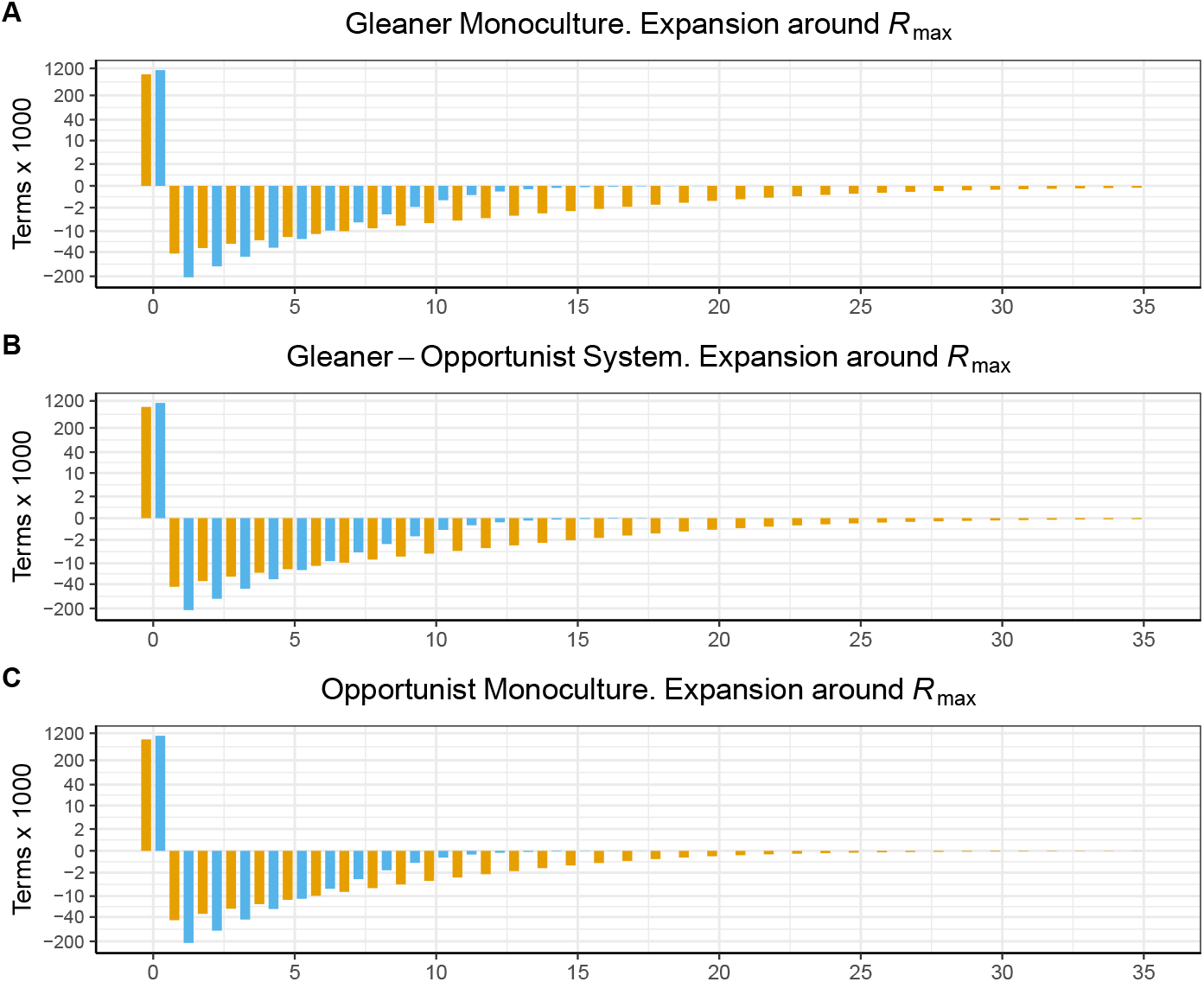
Expansions around *R*_max_ in the gleaner-opportunist system described in the main text. Terms corresponding to the opportunist are shown in blue and corresponding to the gleaner are shown in orange.

**Figure S14:**
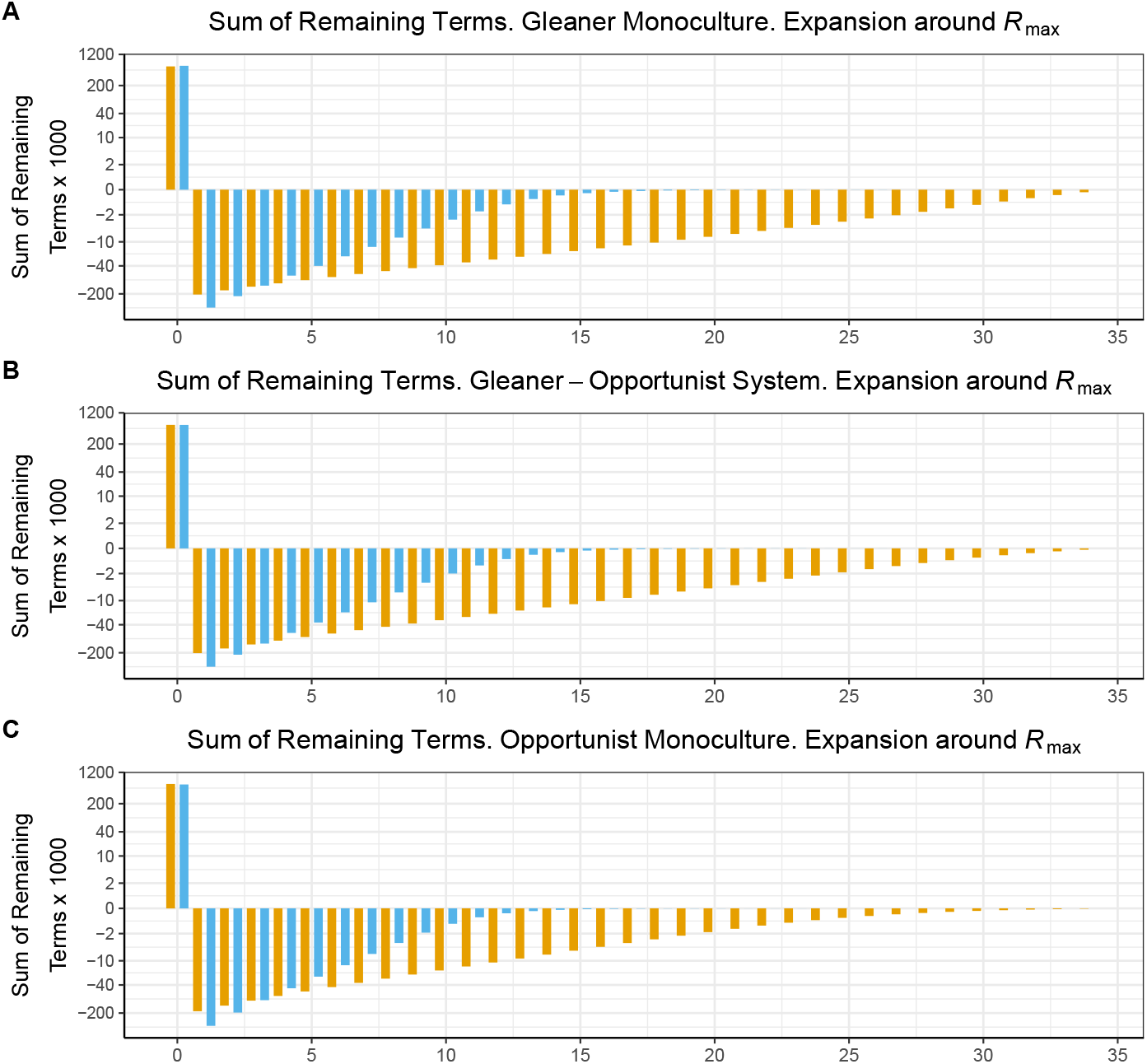
Remainders in the expansions around *R*_max_ in the gleaner-opportunist system described in the main text. Terms corresponding to the opportunist are shown in blue and corresponding to the gleaner are shown in orange.

**Figure S15:**
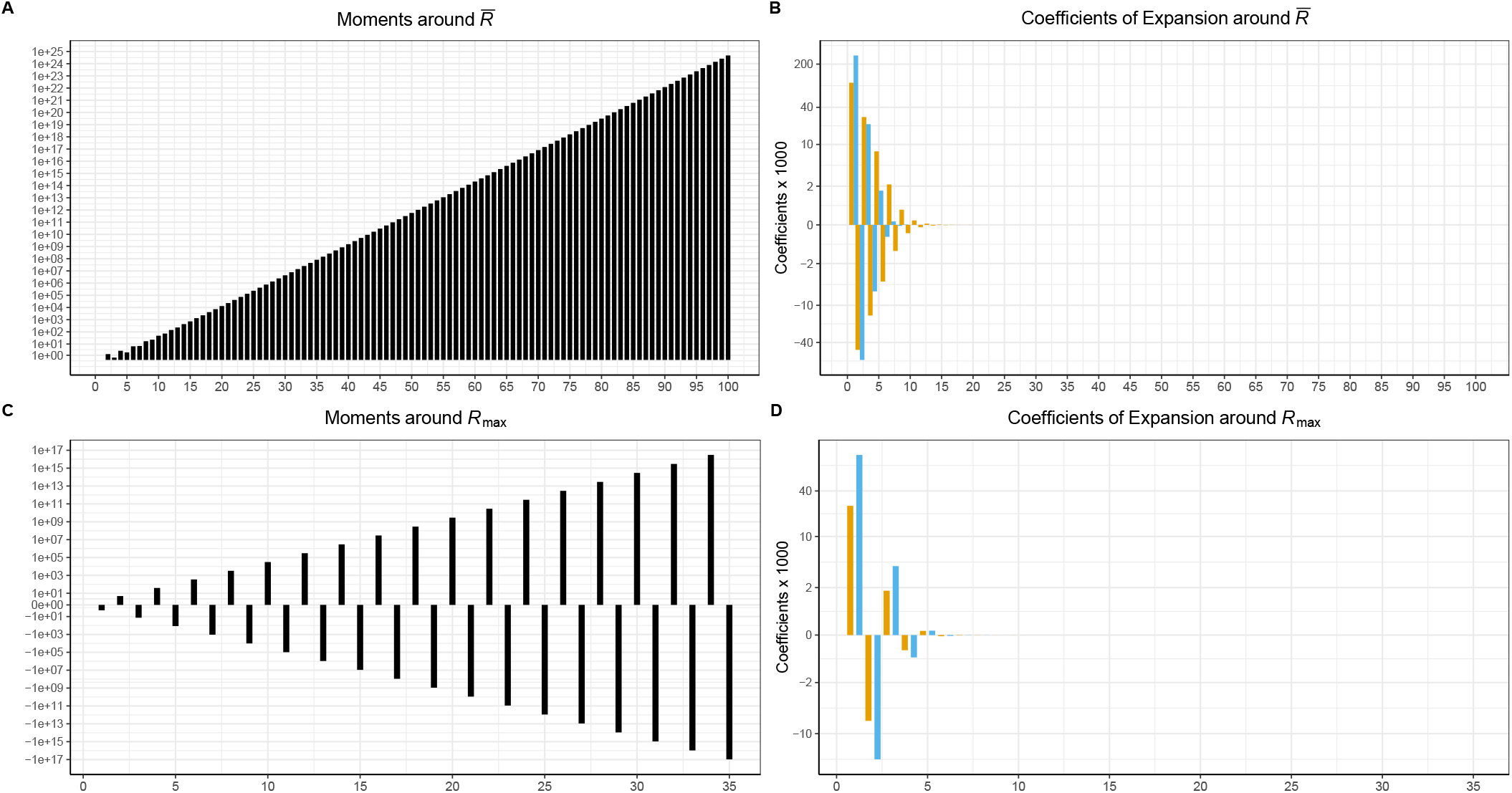
Moments and coefficients in the gleaner-opportunist system described in the main text. Terms corresponding to the opportunist are shown in blue and corresponding to the gleaner are shown in orange.

**Figure S16:**
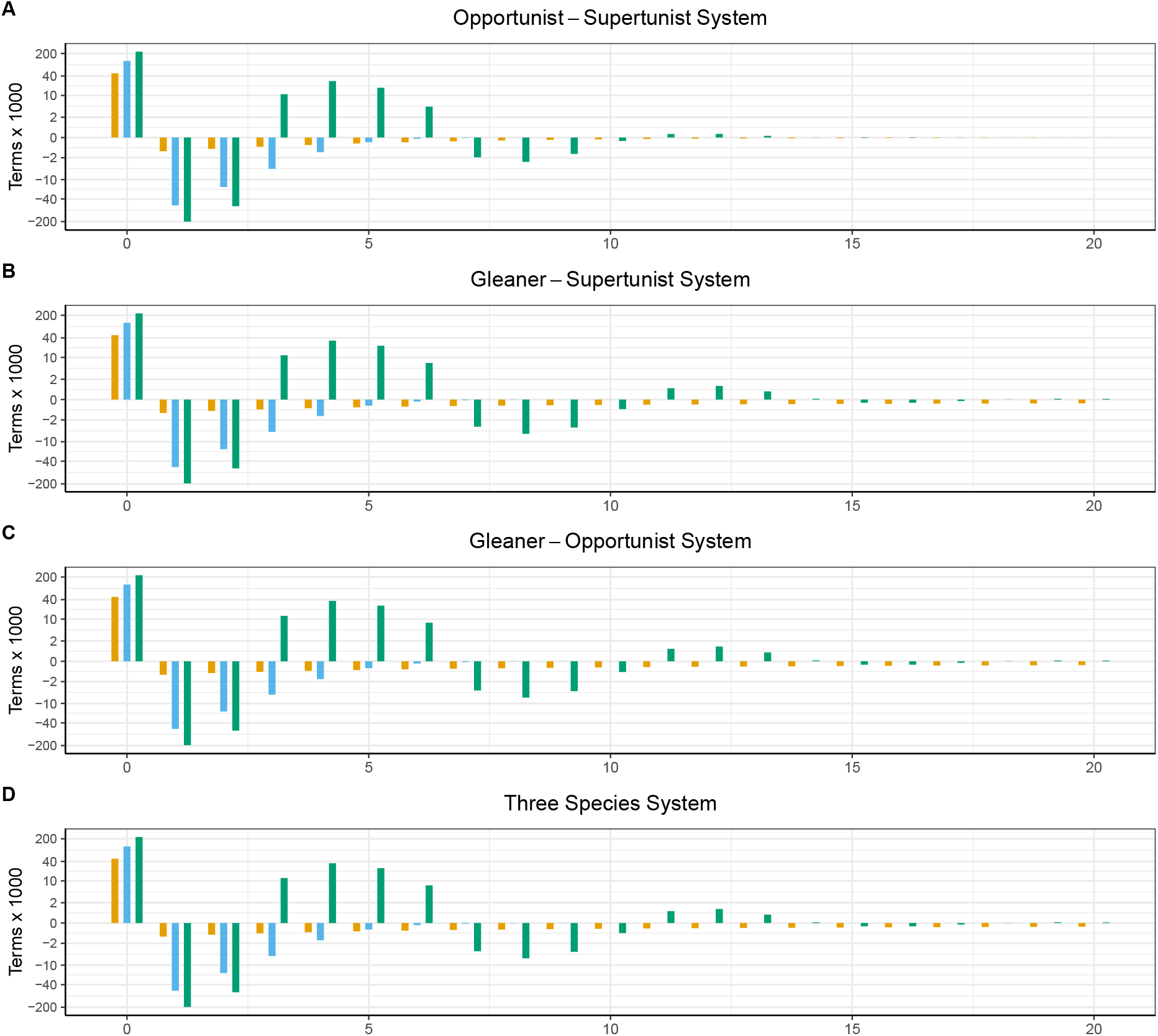
Expansions in gleaner-opportunist-supertunist system described in the main text. Terms corresponding to the opportunist are shown in blue, those corresponding to the gleaner are in orange, and those corresponding to the supertunist are green.

**Figure S17:**
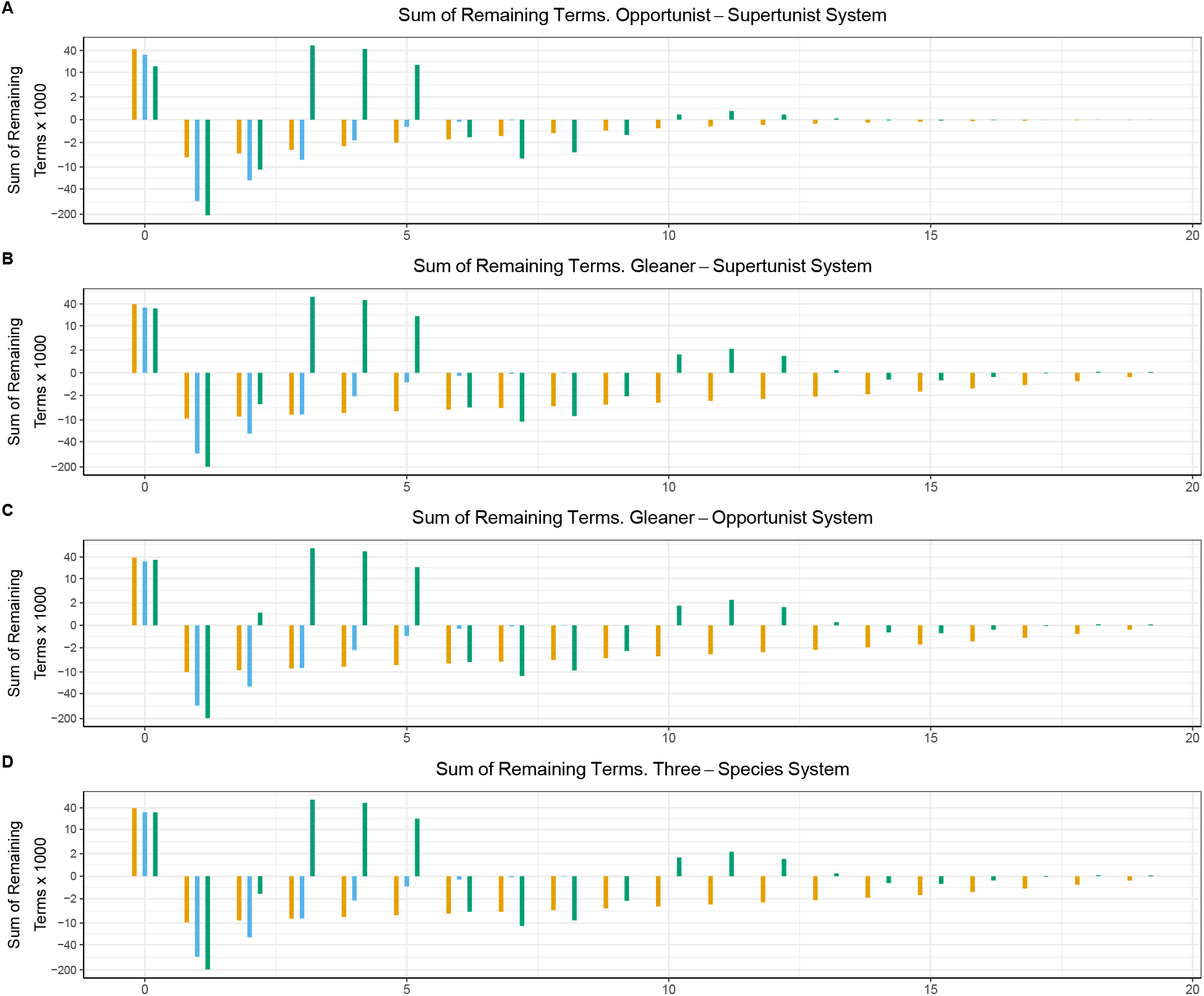
Remainders in gleaner-opportunist-supertunist system described in the main text. Terms corresponding to the opportunist are shown in blue, those corresponding to the gleaner are in orange, and those corresponding to the supertunist are green.

**Figure S18:**
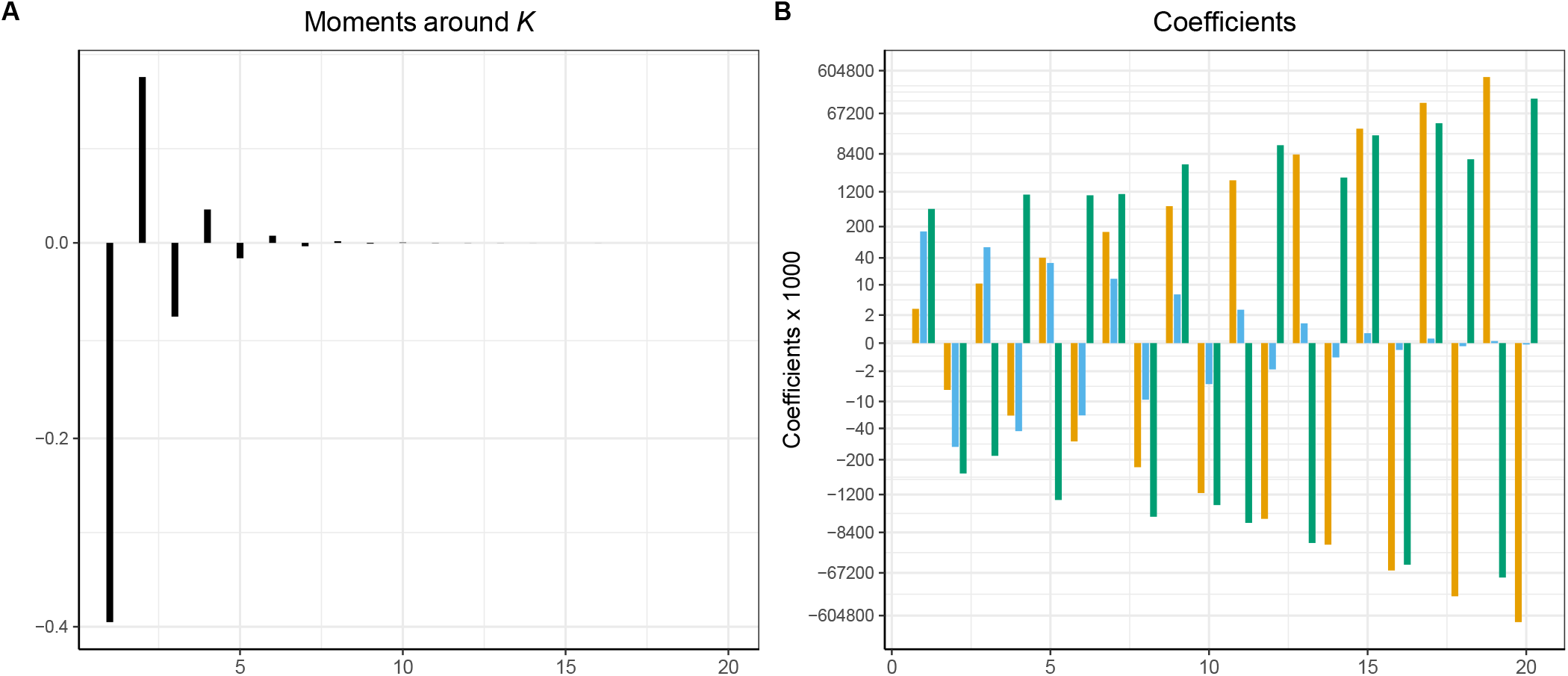
Moments and coefficients in the gleaner-opportunist-supertunist system described in the main text. Terms corresponding to the opportunist are shown in blue, those corresponding to the gleaner are in orange, and those corresponding to the supertunist are green.

**Figure S19:**
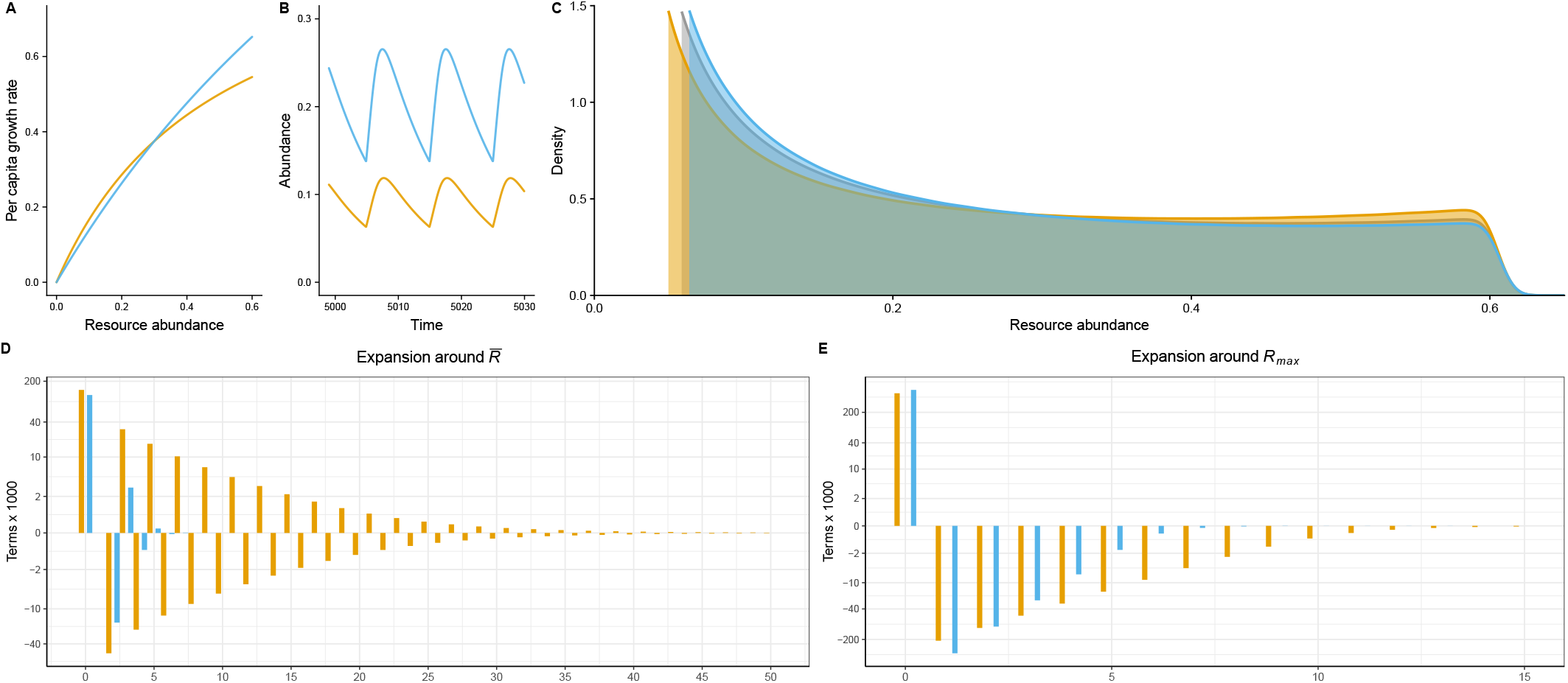
The gleaner-opportunist system with pulsed resource supply (see *S9 A gleaner-opportunist system with discrete resource supply*). **A**. Per capita growth rates, gleanerin orange, opportunist in blue. **B**. Simulation results showing steady-state population dynamics. **C**. Kernel density estimates for resource abundance through time; orange is in the gleaner monoculture system, blue in the opportunist monoculture system, and grey is in the two-species system. **D**. Terms in the expansions of each consumer’s growth rate around 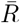 in the two-species system. For visualization, all terms multiplied by 1000. **E**. The expansions of each consumer’s growth rate around *R*_max_ in the two-species system, once again with all terms multiplied by 1000. Note that, both in panel C and panel D, the terms sum to *d*_*i*_ = 0.1.

**Figure S20:**
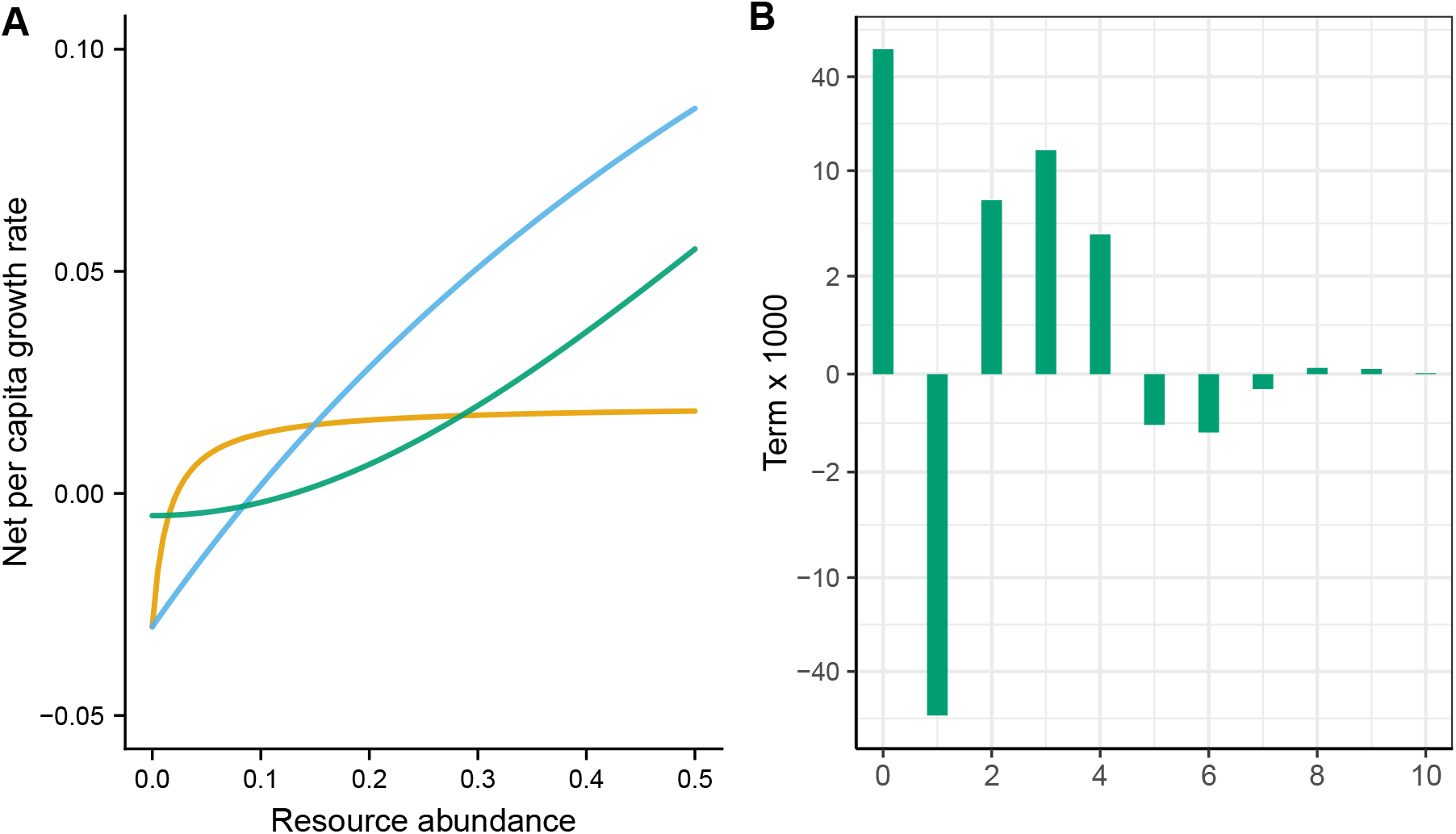
The gleaner-opportunist-survivalist system described in *S10 The gleaner-opportunist-survivalist system*. The parameters are the same as in the gleaner-opportunist-supertunist system with logistic resource supply, except that the survivalist has parameters *r*_3_ = 0.3, *k*_3_ = 1, *l*_3_ = 2, and *d*_3_ = 0.005. **A**. Net per capita growth rates *μ*_*i*_(*R*) − *d*_*i*_ for each consumer. The gleaner is orange, the opportunist blue, and the survivalist green. **B**. The expansion of the survivalist’s average per capita growth rate around *R*_max_, calculated in the three-species system. Note the similarity to the supertunist’s expansion in Fig 4.

**Figure S21:**
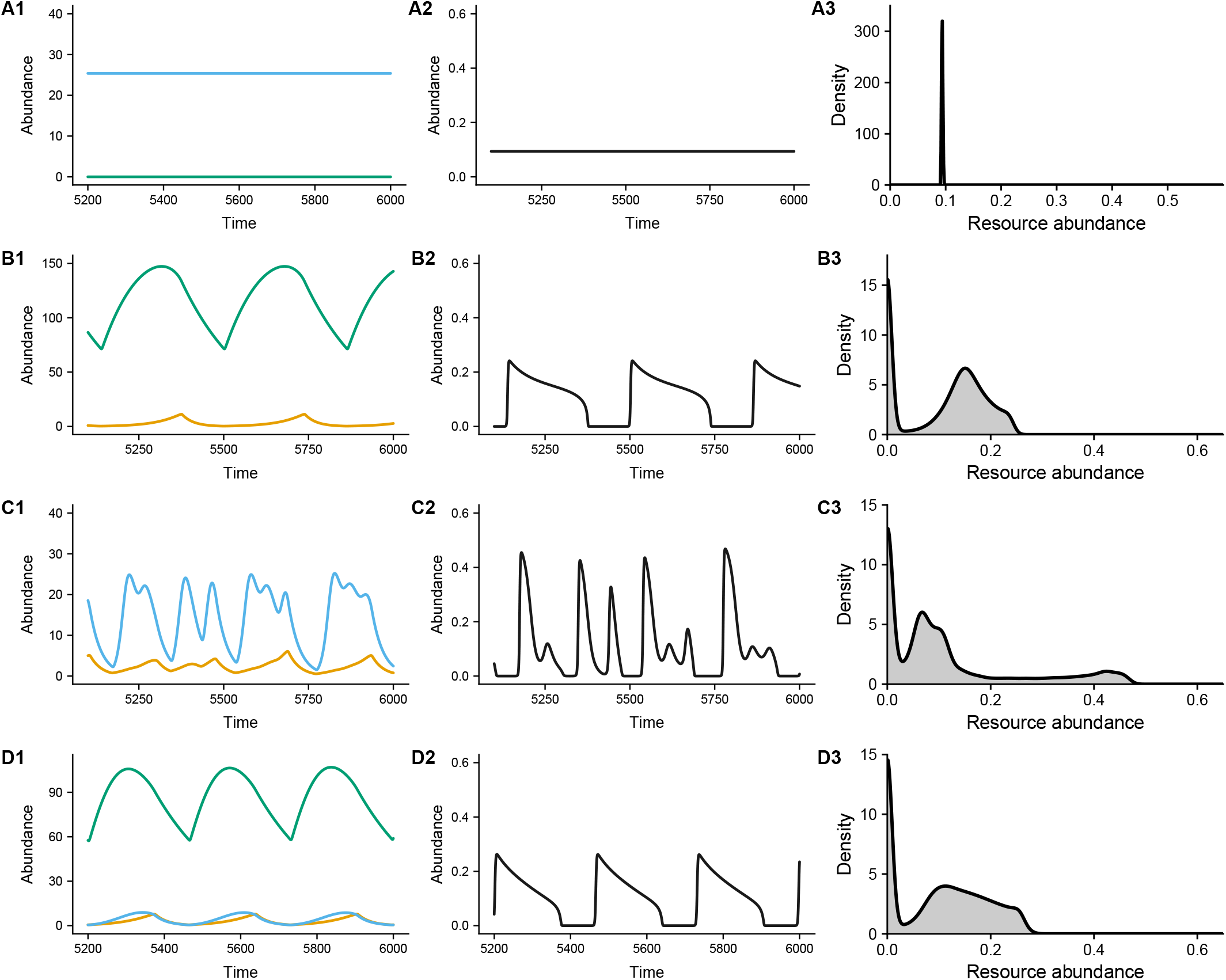
Simulation results for the gleaner-opportunist-survivalist system described in the main text. The opportunist is shown in blue, the gleaner in orange, and the survivalist in green. **A**. The opportunist-survivalist system. **B**. The gleaner-survivalist system. **C**. The gleaner-opportunist system. Note that the gleaner and opportunist in this system are the same as the gleaner and opportunist in the gleaner-opportunist-supertunist system described in the main text. Thus, panel C is identical to the fourth row of Fig 1. **D**. The complete three-species system.

**Figure S22:**
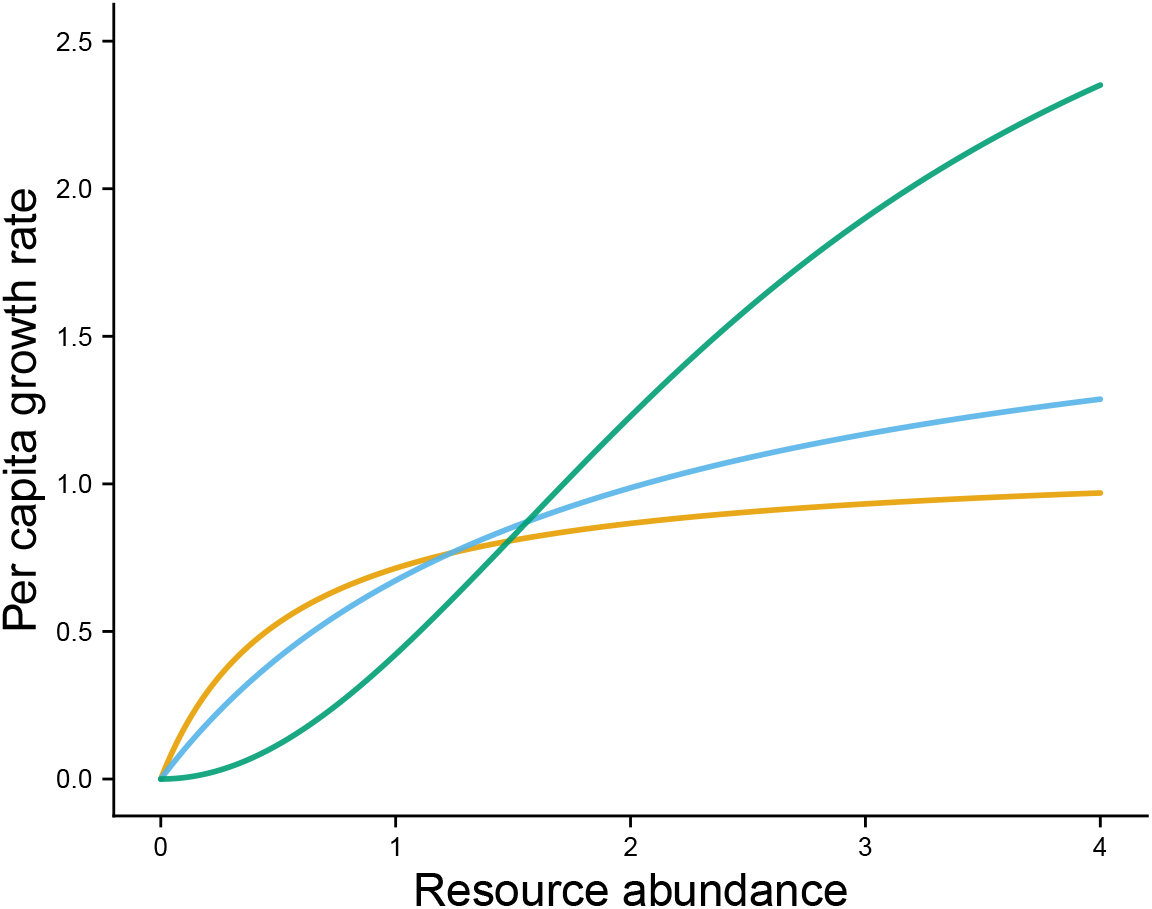
Per capita growth rates for the gleaner-opportunist-supertunist system described in *S11 Community assembly*. The gleaner is orange, the opportunist blue, and the supertunist green.

**Figure S23:**
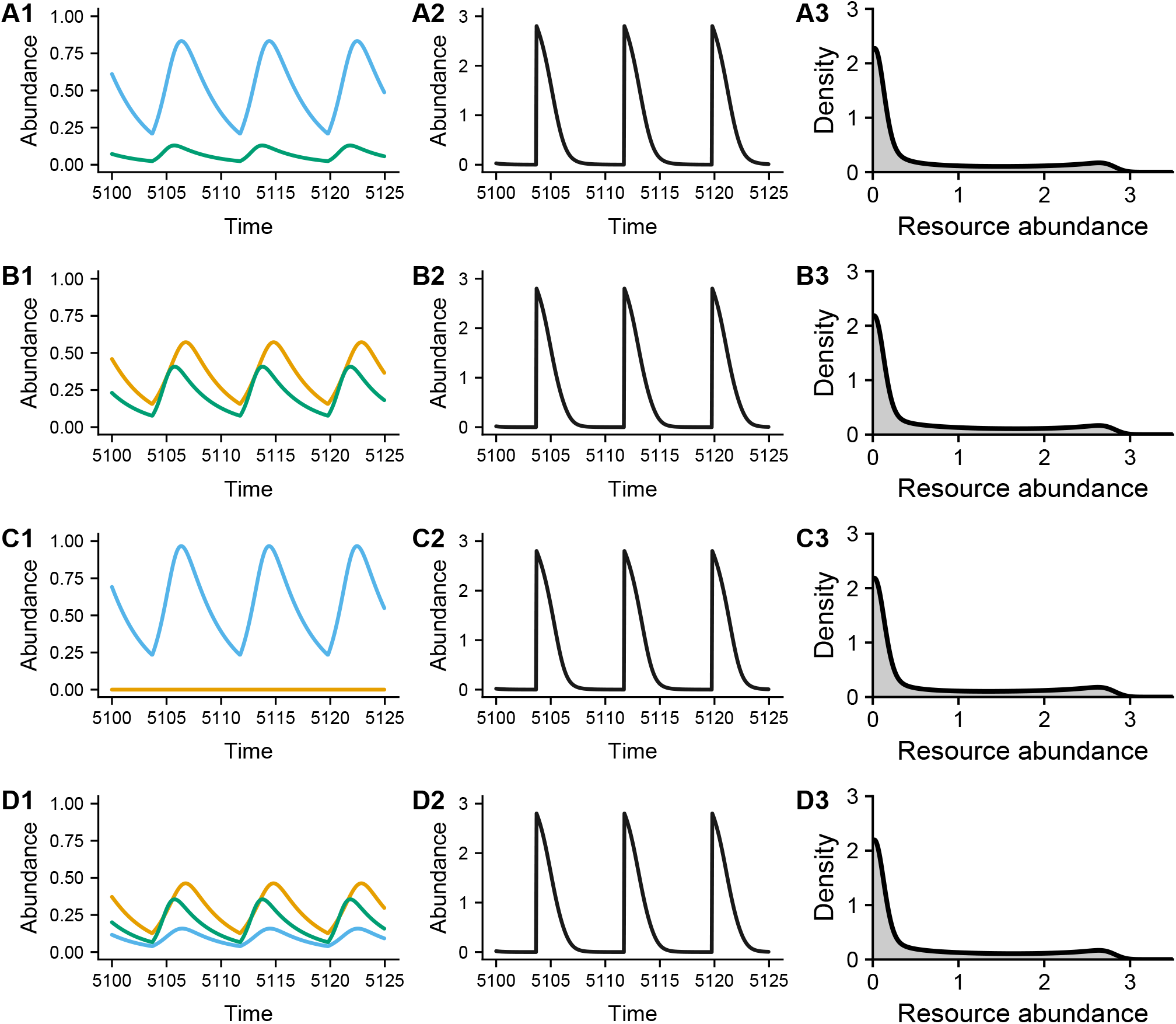
Simulations of the gleaner-opportunist-supertunist system shown in fig S22 with the pulsed resource supply described in *S11 Community assembly*. The opportunist is shown in blue, the gleaner in orange, and the supertunist in green. **A**. The opportunist-supertunist system. **B**. The gleaner-supertunist system. **C**. The gleaner-opportunist system. **D**. The complete three-species system. Note that, with this resource regime, the supertunist acts as a keystone species.

**Figure S24:**
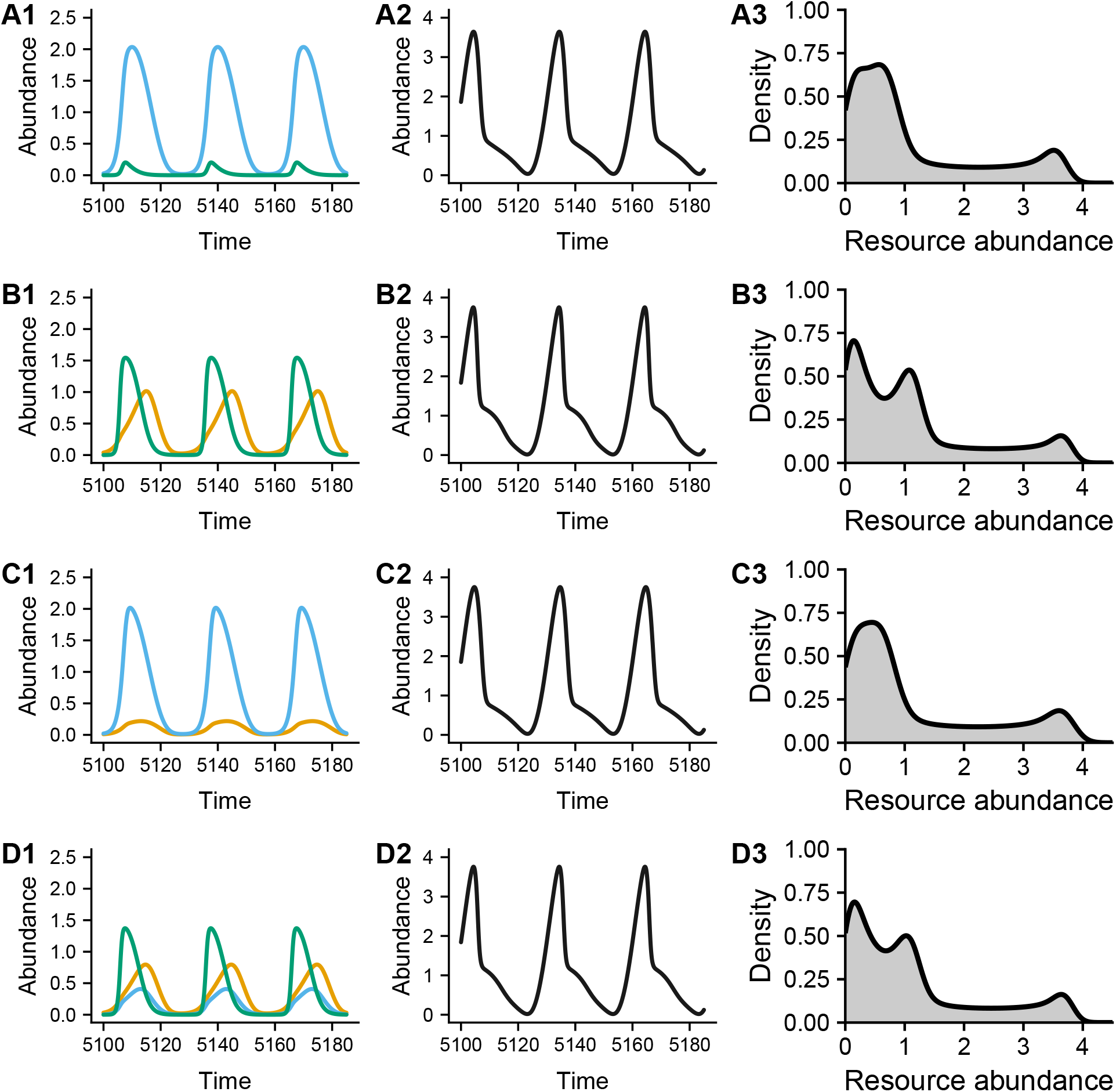
Simulations of the gleaner-opportunist-supertunist system shown in fig S22 with the sinusoidal resource supply described in *S11 Community assembly*. The opportunist is shown in blue, the gleaner in orange, and the supertunist in green. **A**. The opportunist-supertunist system. **B**. The gleaner-supertunist system. **C**. The gleaner-opportunist system. **D**. The complete three-species system. Note that, with this resource regime, the system has no keystone species.

